# “Quasigenus” among *Phycodnaviridae*: A diversity of chlorophyte-infecting viruses in response to a dense algal culture in a high-rate algal pond

**DOI:** 10.1101/2022.12.26.521929

**Authors:** Emily E. Chase, Thomas M. Pitot, Sonia Monteil-Bouchard, Christelle Desnues, Guillaume Blanc

**Affiliations:** Department of Microbiology, University of Tennessee, Knoxville, Tennessee 37996, USA; Department of Biochemistry, Microbiology and Bioinformatics, Laval University, Quebec, Canada; Microbiologie Environnementale Biotechnologie, Institut Méditerranéen d’Océanologie, Campus de Luminy, 163 Avenue de Luminy, 13009 Marseille, France; Institut hospitalo-universitaire (IHU) Méditerranée infection, 19-21 Boulevard Jean Moulin, 13005 Marseille, France

**Keywords:** *Nucleocytoviricota*, polinton-like viruses, virophage, metagenomics, microalgae

## Abstract

This study approaches a high rate algal pond (HRAP) culture by metagenomic sequencing of the viral DNA fraction, this includes the so-called giant virus fraction (phylum *Nucleocytoviricota*), with the goal of revealing viruses coexisting within an intensified algal culture. A wealth of interesting novel viruses is revealed, including members of *Nucleocytoviricota, Lavidaviridae*, and polinton-like viruses, which are taxa containing previously characterized algal viruses. Our sequencing results are coupled with a virus targeted qPCR study and 18S rDNA metabarcoding to elucidate potential virus-host interactions. Several species of green algae are identified (Chlorophyta), likely representing the alternating dominant populations during the year of study. Finally, we observe a bloom of viral diversity within the family *Phycodnaviridae* (*Nucleocytoviricota*), including highly related but non-identical genotypes, appearing in the HRAP in September and October 2018. This bloom is most likely the cause of a mass mortality event of the cultured algae that occurred during these same months. We hypothesize that these related *Phycodnaviridae* lineages selectively infect different strains of the same algal species of the Genus *Picochlorum* that have been identified in the HRAP by metabarcoding and coined this phenomenon a “quasigenus” by analogy to the RNA virus quasispecies concept.

## Introduction

The importance of viruses in aquatic systems is well documented from nutrient cycling and algal bloom control in the open ocean (Suttle and Wilhelm, 1999; Biggs et al., 2021), to modulating microbial communities in the deepest ocean (Jian et al., 2021), and throughout the open ocean water column (Luo et al., 2020). In relation to microalgae, specific viral groups of interest include members of Phylum *Nucleocytoviricota* or “giant viruses”, virophages (*Lavidaviridae*), and polinton-like viruses. Family *Phycodnaviridae* and family *Mimiviridae* are two taxa within *Nucleocytoviricota* known to infect microalgae (Wilson et al., 2009; Claverie and Abergel, 2018). Family *Lavidaviridae* are a group of viruses infecting *Nucleocytoviricota* in a tripartite/co-infection system between a giant virus and its host (Fischer, 2021). Previous work on family *Lavidaviridae* has given evidence for algal *Nucleocytoviricota* hosts of these tripartite systems (Yau et al., 2011), and recently characterised this interaction *in situ* with a *Chlorella* host (Sheng et al., 2022). Despite evidence of a virophage benefiting the cellular host indirectly by reducing *Nucleocytoviricota* production in co-infection (La Scola et al., 2008), and regulating host-virus dynamics (Yau et al., 2011), much is still unknown about family *Lavidaviridae*. Polintons (or Mavericks), are relatively large (up to 40 kb) DNA transposons and have been found in a substantial diversity of multicellular and unicellular species, including protists, fungi, beetles, fish, chicken, etc. (Kapitonov and Jurka, 2006). In 2015, a 31 kb DNA virus (TsV-N1) similar in structure to polintons and with which it shares a number of phylogenetically related genes, was identified as viral particles infecting the microalgae *Tetraselmis striata* (Pagarete et al., 2015). This confirmed that some polinton-like viruses (PLVs) behaved as genuine viruses (*i*.*e*., having infectious virions) and not just transposable elements (Koonin and Krupovic, 2017). Notably, the PLVs have some differences from polintons, including the absence of specific genes (see (Yutin et al., 2015)). Recent metagenomic work concluded that there are at least eight major groups of PLVs and also uncovered many associated with microalgae (Bellas and Sommaruga, 2021). Interestingly, members of family *Lavidaviridae* can possess similarities to PLVs (Koonin and Krupovic, 2017) and PLVs also have potential as a co-infection with extended *Mimiviridae* species (Yutin et al., 2015), therefore could contribute to viral-host dynamics in a tripartite system as well.

Given the significance of viruses in major aquatic systems our study focuses on a specific system that is relatively unexplored until now; the viral community of an industrial high-rate algal pond (HRAP). The said HRAP hosts a non-specific polyculture of microalgae sourced from seawater of the Mediterranean Sea. Our objectives were to understand the diversity of microalgal viruses in the system alongside their dynamics during a culturing period. More specifically, the HRAP environment permitted a dense microalgal culture not unlike an algal bloom of a natural aquatic environment, consequently providing an opportunity to investigate a bloom-like situation and the viral community during this period. Each of the aforementioned virus groups (*i*.*e*., *Nucleocytoviricota*; family *Phycodnaviridae* and family *Mimiviridae*, family *Lavidaviridae*, and PLVs) were investigated within the microalgae culturing HRAP of this study to gain an understanding of their role in a dense culture that we suggest is analogous to a natural microalgal bloom.

## Materials and methods

### Sampling, filtration, nucleic acid extraction, and next-generation sequencing of HRAP sampled water

Water sampling and HRAP characteristics are as described in a previous publication (Chase et al., 2021). Samples were processed (*e*.*g*., centrifuged, filtered, etc.) in the same manner to produce an “ultravirome” (UV) of “small” DNA viruses (*i*.*e*., <0.2µm). As an attempt to isolate and subsequently sequence *Nucleocytoviricota*, fluorescence activated cell sorting (FACS) was carried out in the size fraction 0.2 μm – 1.2 μm labelled using SYBR-Green and analyzed according to the side scatter and FITC fluorescence parameters. The size fraction was obtained by filtering samples through a 1.2 μm pore filter (Sartorius ref: 17593), then concentrated 25× on a tangential flow filtration column with a nominal pore size of 0.2 μm (Microkros ref: C02-P20U-05-N). Four gated populations, “megaviromes” (MV), corresponding to areas of the SSC-FITC plot (where giant viruses are generally observed (Khalil et al., 2016)) were sorted in each sample (**Figure S1**). Note that some bacterial populations may emerge in the same area (i.e., gated population) and thus are considered “contaminants” in the MVs. The number of isolated particles (events) per population ranged between 89 000 and 500 000. Lysis was carried out as detailed in (Chase et al., 2021), and nucleic acid was extracted using EZ1 Advanced XL extraction with a Virus Card (QIAGEN). Six of the 20 FACS populations did not generate sufficient DNA and were not subsequently sequenced. Library preparation and paired-end 2 × 250 bp Illumina MiSeq sequencing of the UVs were done as described in a previous publication (Chase et al., 2021). FACS population were amplified using Ready-To-Go Genomiphi V3 kit (GE Healthcare), followed by purification and Illumina HiSeq sequencing (paired-end 2 × 151 bp) performed by DOE Joint Genome Institute (JGI).

### Quality control and contig assembly of metagenomes and putative taxonomic assignments

Quality control steps with raw reads were as described in previous work (Chase et al., 2021). Assemblies of UVs and MVs were done with SPAdes version 3.15.0 using the metaviral algorithm and the single cell algorithm respectively (Antipov et al., 2020). Only contigs over 2000 bp were kept for further analysis. Assembly statistics were done using metaQUAST version 5.1 (Mikheenko et al., 2016) and are reported in **Table S1**. Taxonomic assignments were given by an “open reading frames (ORFs) voting” procedure. To fulfill this, ORFs greater than 100 codons were extracted from each contig in the UVs and MVs, aligned against TrEMBL (Boeckmann et al., 2003) using MMSEQ (Steinegger and Söding, 2017) with e-value < 10^−05^ and the taxonomic information of the ORF best hits were recorded. Contig taxonomic classification was done using a majority rule criterion over the ORF best hits. On average, 92.7% of the contigs were classified within the UVs and 97.1% within the MV contigs. Overall, 3335 contigs were specifically assigned to viruses (prokaryotic or eukaryotic) from which 36,525 ORFs were extracted. Further ORFs functional annotation was performed by protein alignment against the SWISS-PROT (Boutet et al., 2007) database using MMSEQS. We used the eggnog-mapper (Cantalapiedra et al., 2021) for assignment to COG (Tatusov et al., 2000) and KEGG (Kanehisa et al., 2016) functional categories.

### Identifying potential microalgae hosts using 18S rDNA metabarcoding of HRAP water

To characterize the diversity of eukaryotic microorganisms by metabarcoding, DNA was extracted from 0.2 μm filtered water and the V4 variable region of the 18S ribosomal RNA gene was amplified using eukaryote-specific universal primers (forward CCAGCASCYGCGGTAATTCC and reverse ACTTTCGTTCTTGATYRA; (Guillou et al., 2013)). The obtained amplicons were then subcontracted by the Genotoul facility (Toulouse; get.genotoul.fr) which performed indexing and Illumina MiSeq sequencing. The sense and antisense sequences obtained were assembled and then clustered as ASVs (amplicon sequence variants) and identified at different taxonomic levels using the PR2 database version 4.12.0 (Guillou et al., 2013). These analyses were performed under R software version 3.6.2 using the *DADA2* package (Callahan et al., 2016).

### Phylogenetic construction of Phylum Nucleocytoviricota, PLVs, and family Lavidaviridae relationships

A major capsid protein (MCP), DNA polymerase B (PolB), and ATPase tree were produced using publicly available data for *Nucleocytoviricota* using an alignment from MAFFT v.7 (Katoh and Standley, 2013), and FastTree (Price et al., 2009) with default settings. Additionally, a PLV tree (MCP based using publicly available data) was composed also using FastTree with default settings. MCP sequences were downloaded from National Centre for Biotechnology Information (NCBI) for family *Lavidaviridae*. Corresponding sequences were also aligned using MAFFT v.7 and the *Lavidaviridae* tree was produced using FastTree with default settings. All trees were run with 1000 bootstrap replicates. In all cases putative sequences from relevant groups recovered from the HRAP were included in the trees.

### Dynamics of putative viral targets by qPCR

The selection of putative viruses for targeting, processing of raw water samples for qPCR, and the qPCR reaction setup were conducted in the same way as described in previous work (Chase et al., 2021) without the need for cDNA construction. Briefly, Primer3 (Untergasser et al., 2012) and PerlPrimer v.1.2.3 (Marshall, 2004) were used to produce efficient primers *in silico* before *in vitro* testing. The number of reaction cycles (total of 45) minus the Cq value were reported as the “inverted Cq”.

### Using correlation among viruses and potential hosts to uncover possible host-virus interactions

Principle component analyses (PCA) were performed and visualised on a combined dataset of standardized metabarcoding (18S) and qPCR results using R package *factoextra*, using time points where both metabarcoding and qPCR results were available. Similarity profile analysis (SIMPROF) hierarchical clustering at α=0.1 was conducted using R package *clustsig*, and correlation networks were visualised using Cytoscape v3.9.1 (Smoot et al., 2011). Publicly available sequenced alga of potential hosts (*e*.*g*., Picochlorum *spp*.) were checked for recent *Nucleocytoviricota* viral insertions using ViralRecall (Aylward and Moniruzzaman, 2021).

## Results and Discussion

### Metagenomic sequencing data and biodiversity of putative DNA viruses in the HRAP

In total five ultraviromes (UV) from different months of 2018 (*i*.*e*., April [04.17], May [05.17], July [07.05], September [09.11], and October [10.23]) were sequenced, and 14 megavirome (MV) populations based on FACS gated populations (see **Table S1**). Following genome assembly, the number of contigs greater than 2 kb ranged from 74 to 1,968 for the UV samples, and 180 to 10,045 for the MVs, with the average number per sample being 1221 and 3995 for the UV and MV respectively. The GC content in the UV ranged from 47% to 54%, whereas in the MV it ranged from 38% to 53%. Bacterial sequences were very abundant in all viromes (**Figure S2**). Regardless of potential contamination issues that are common in viral metagenomics (Jurasz et al., 2021), this result is not surprising for FAC-sorted MVs, as the size fraction of sorted particles includes that of small bacteria. For UVs, this could be due to leaks in the filtration or inefficient digestion of free DNA (lysed bacteria) present in the sample. Nevertheless, virus contigs still represented between 0.4% (05.17.P4) to 39% (07.05.UV) of the cumulated contig length. Eukaryotic virus sequences were generally dominant over prokaryotic virus sequences in MVs (the mean proportion was 2.5% for eukaryotic viruses versus 0.8% for prokaryotic viruses) while this trend was opposite in the UVs (mean frequency was 18.1% versus 0.6%, respectively).

Using metagenomics, we employed two approaches to uncover DNA viruses of the HRAP, permitting the observation of small DNA viruses within a UV, and large DNA viruses (*e*.*g*., *Nucleocytoviricota*) through FACS populations and further processing (MV). The UVs are primarily composed of phages (*e*.*g*., *Siphoviridae, Podoviridae*, and *Myoviridae*), and unclassified prokaryotic viruses (**Figure 1 A**). Polinton-like viruses were also uncovered in the DNA UVs and MVs (appear in the “unclassified virus” category; see also phylogeny in **Figure S3**) and are discussed more below. Of special importance to our study, is our effort to retrieve *Nucleocytoviricota* by FACS methods, and we were able to recover a viral diversity community composition different than the UV fraction (**Figure 1 B**). Recent work (Palermo et al., 2021) on metagenomic sample filtration methods points out the almost definite loss of viral community taxonomic information in metagenomic studies where larger (> 0.45 μm pore size) water fractions are removed and left un-sequenced. No doubt, sequencing this large viral fraction permitted higher recovery of *Nucleocytoviricota* compared to the UV method. A putative member of family *Lavidaviridae* (a virophage) was detected and does not group closely with a *Lavidaviridae* known to infect a *Chlorella* species (**Figure S4**). As stated previously, virophages are found in association with algae infecting *Nucleocytoviricota*, and were recovered in our MV samples (**Figure 1 B**). This result will be explored in more depth.

**Figure 1.**
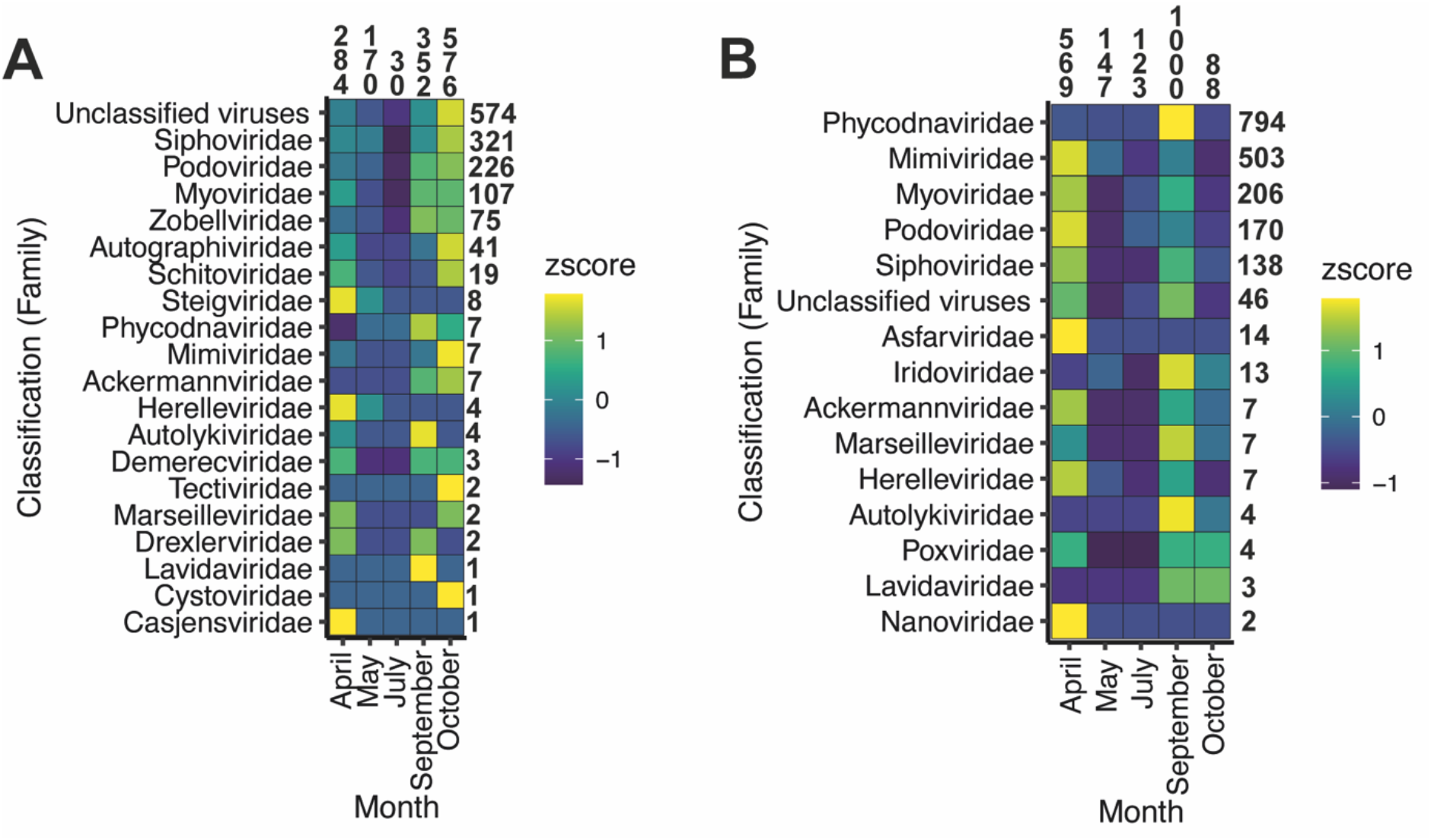
Proportion (by z-score) of viral taxonomic assignments to assembled (**A**) DNA ultravirome contigs by each metagenomic sample in 2018, and (**B**) proportion of viral taxonomic assignments to assembled metagenomes produced from FACS population, grouped together by sampling month. Please note (**B**) is only to showcase the diversity and is not a quantitative assessment as some sample months produced more FACS populations of interest, and therefore were attributed more sequencing. Additionally, these classifications are resulting from our “ORF voting” procedure. Total numbers of contigs attributed to each family are shown on the y-axis, and total number of contigs attributed to each month are shown on the x-axis.

### Genetic potential of viral communities

Of 21,896 ORFs extracted from eukaryotic virus contigs, 5407 (25%) had a significant match in sequence databases, and 811 could be assigned to COG (Cluster of Orthologous Groups of proteins) functional categories (**Table S2**). Also, 198 ORFs were identified as encoding an MCP. For prokaryotic viruses contigs, 6586 (45%) of the 14,629 extracted ORFs had a match in sequence databases, of which 154 were against a capsid protein and 728 were assigned to a COG functional category. Contigs of prokaryotic viruses were found to encode a higher proportion of proteins involved in DNA replication and repair, and in metabolisms of membranes and cell walls (**Figure S5**). One possible reason is that these viruses were mainly found in the UV fractions and therefore, likely had smaller genomes carrying genetic information more narrowly focused on essential viral functions, *i*.*e*., replication and entry into the cell. Conversely, eukaryotic viruses, which were more highly represented in the MV fractions, encoded a greater proportion of proteins involved in accessory metabolic functions, including nucleotide metabolism, protein translation and post-translation modification.

Genes of viral contigs that had significant matches in cellular organisms but not in viruses were further examined. These genes are probably derived from horizontal transfers from cellular hosts. They represent candidate auxiliary metabolic genes (AMGs) that have not yet been observed in viruses or whose sequence divergence is such that the homology relationship with another virus is no longer detectable by protein alignment.

Eukaryotic virus contigs carried 99 of these potential AMGs, 62 of which had an unknown function. Among the genes to which a function could be attributed, several were located on contigs generated from the 07.05.P3 population and encoded remarkable functions including a neutral sphingomyelinase which is a key enzyme in sphingolipid metabolism and is implicated in stress-induced signaling pathways (Choezom and Gross, 2022). Another gene encoding a myosin motor domain-containing protein suggests that the corresponding virus might use actin-based movements in the infection process. Contigs from different populations of September (09.11) also encoded a variety of interesting proteins never yet observed in eukaryotic viruses. They include homologues of heme oxygenase and Phycocyanobilin:ferredoxin oxidoreductase which are both involved in synthesis of bilins which play multiple role in chlorophyll metabolism and signal transduction (Zhang et al., 2018), a CP12-like protein involved in regulation of the Calvin cycle responsible for CO_2_ assimilation (Gontero and Maberly, 2012), a RNA polymerase subunit Rpb5, a phosphatidylserine decarboxylase involved in the synthesis of the abundant phospholipid phosphatidylethanolamine of mitochondrial membranes (Nerlich et al., 2007), a clavaminate synthase-like proteins potentially involved in the biosynthesis of clavulanic acid and other 5S clavams (Tahlan et al., 2004), which have antibacterial and antifungal activities, a non-histone chromosomal MC1 family protein whose archeal homologue is used for DNA packaging (De Vuyst et al., 2005), and a VIT1/CCC1 iron transporter family protein. We also found components of restriction modification (RM) system, including several endonucleases and DNA methyltransferases yet unseen in viruses. The presence of RM systems is unsurprising, given that they are found in members of *Nucleocytoviricota* (Filée, 2018), and more specifically have been documented in *Phycodnaviridae* (chloroviruses) (Coy et al., 2020). As previously summarized (Jeudy et al., 2020), nucleases within giant virus genomes could assist in the degradation of host DNA to permit recycling into viral particles, or inhibit host gene expression (Agarkova et al., 2006). Prokaryotic virus contigs encoded 242 proteins that match homologs in cellular organisms but not in viruses, of which most (209) had an unknown function. Interesting functions were identified, including a ABC transporter efflux protein encoded by a prokaryotic virus from the 09.11.P4 (MV) population, a N-acetylmuramoyl-L-alanine amidase (09.11.UV) involved in cleavage of cell-wall glycopetides, a helix-turn-helix (HTH) transcription regulator (05.17.UV), a non-viral-type sialidase (10.23.UV) involved in the degradation of extracellular mucin, a RNA polymerase sigma factor (04.17.UV) and a trehalose utilization ThuA protein. Taken together, these results suggest that HRAP viruses encode a specific subset of regulatory and/or enzymatic proteins that differs from the proteomes of model viruses that serve as a template for comparative studies.

### Biodiversity of potential alga hosts in the HRAP

When considering only the top ten classifications of V4-18S ASVs in each sample month (**Figure 2 A**) the most abundant taxa are class *Trebouxiophyceae* (primarily family *Chlorellales*), and for the scope of this paper will be the focus. It is worth noting, the lack of diatoms is most likely due to the HRAP culture lacking in biogenic silica (*i*.*e*., no silica was added). Of specific interest to our study, several groups of unicellular microalgae are present (**Figure 2 A** and **B**). Including aforementioned family *Chlorellales*, alongside other members of phylum Chlorophyta; *Chlorodendraceae, Chlamydomonadales, Sphaeropleales, Pyramimonadales, Ulvophyceae*, and *Mamiellaceae* in the HRAP during the 2018 culture samples (highlighted in **Figure 2 B**). Given previously established associations with DNA viruses to Phylum Chlorophyta (Etten et al., 2020; Short et al., 2020), we focused mostly on these microalgae and not additional unicellular plankton identified in the HRAP by metabarcoding, as these species do not appear in substantial amounts (based on ASVs) in 2018 alongside our metagenomic study or present evidence of relationships with the putative viruses tracked in our study.

**Figure 2.**
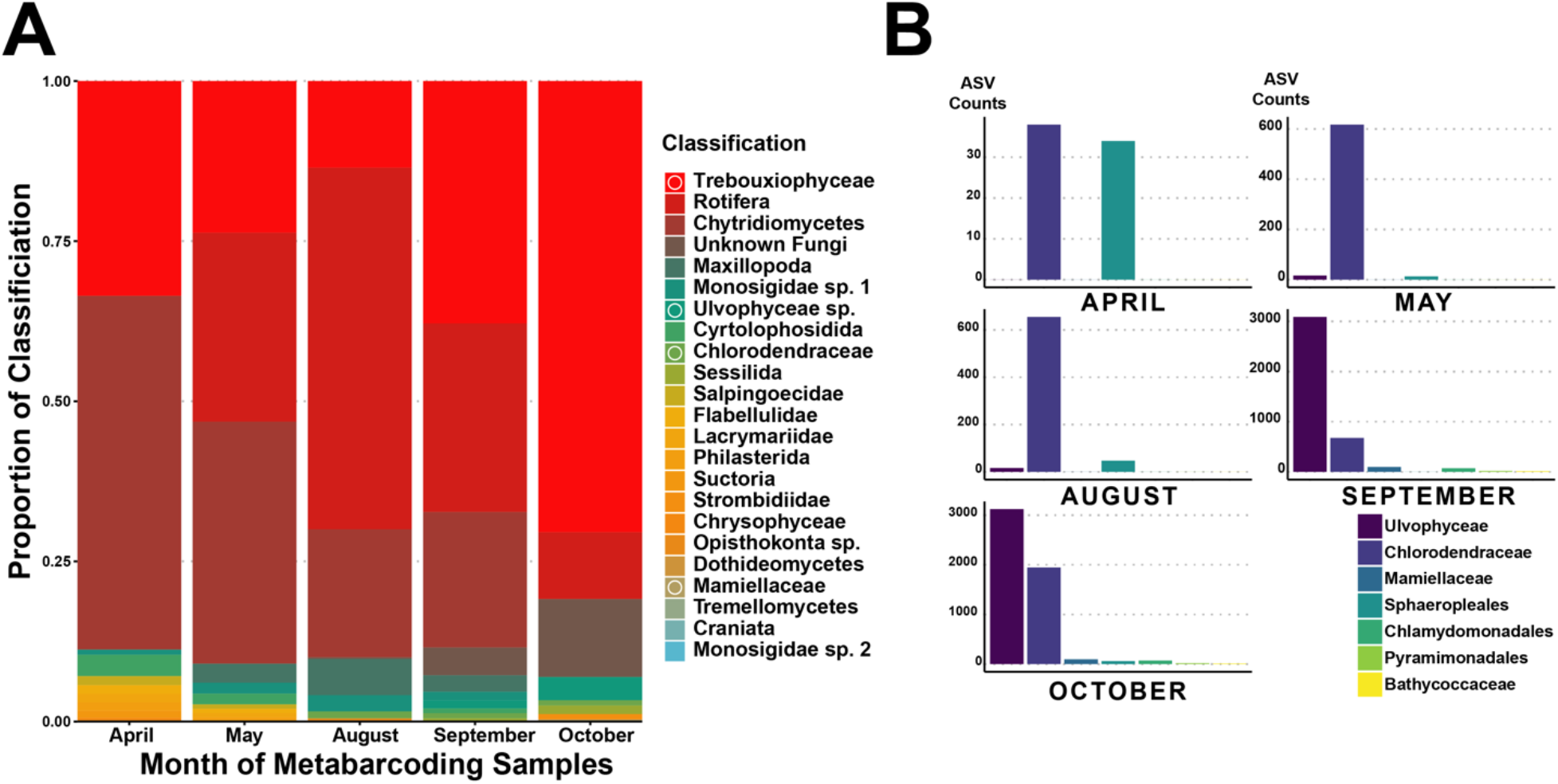
Classification of HRAP eukaryotes by **(A)** proportion of amplicon sequence variants (ASVs) assigned to taxonomy of 18S metabarcoding data for each month of 2018 that also has metagenomic sampling, ASV from each month were summarised. Members of Phylum Chlorophyta are indicated on the legend with a circle. Only the top ten ASV classifications for each month are represented in the schematic. Additionally, **(B)** ASV counts of all Chlorophyta (excluding the abundant Class Trebouxiophyceae), this is not limited to the top ten ASV classifications per month.

### Predicting potential microalgal hosts and their viruses

Using correlation of microalgal species and viral population presence we can try to resolve potential host and virus relationships (Hingamp et al., 2013; Roux et al., 2017). As the approach is simply observing correlations it cannot account for ecological nuances of host-virus relationships, namely it assumes that overlapping host and virus presence and their relative abundances are an indicator of infection. With this understanding we interpreted our results as a preliminary suggestion of possible host-virus relationships in the context of known viral ecology. Correlation analyses were inconclusive for virus-host relationships outside of Chlorophyta alga, and consequently we focused solely on this group.

Principal component analysis (PCA) with members of familys *Mimiviridae, Phycodnaviridae*, and *Lavidaviridae* (sequenced within MVs) showed a strong positive correlation (*i*.*e*., higher covariance) among *Picochlorum* and several members of these viruses (**Figure S6 A**), including a putative phycodnavirus that also had a strong correlation with the detected member of family *Lavidaviridae* (*i*.*e*., the putative virophage). A strong correlation was shown between a member of family *Mimiviridae* (M1) and the alga group *Dictyosphaerium* (**Figure S6 A**). These relationships are primarily associated with September and October (2018) sampling. Although much less supported by these principal components, there was an additional association between a member of family *Mimiviridae* (M2) and an unknown species from the order *Chlorellales*. The PCA for polinton-like viruses (PLVs) does not show a close correlation for some of the putative viruses (**Figure S6 B**). Some PLVs are associated with genera *Dictyosphaerium* and *Mantoniella*, and to some extent the genus *Picochlorum*. However, these variables contribute much less to the ordination than the examples highlighted within the *Nucleocytoviricota* and *Lavidaviridae* PCA (**Figure S5 A**). Hierarchical clustering indicates a significant grouping among previously described MV viruses and hosts (**Figure S7; b, d**, and e). Additionally, PLVs and their associated PCA suggested hosts also produce significant groupings, although with slightly weaker correlations (although still significant) in most cases (**Figure S8**) in relation to *Nucleocytoviricota* and *Lavidaviridae* groupings. Correlations among both the MVs and PLVs with genus *Picochlorum* reflect an apparent bloom of *Picochlorum* in the area the HRAP is stationed during these months, however in comparison to the PLVs the correlation among 11 *Phycodnaviridae*, one *Mimiviridae*, and the putative virophage are specifically much stronger (**Figure S7** b). In the context of viral ecology, it seems more likely that correlations found among *Nucleocytoviricota* and *Lavidaviridae* could imply potential hosts more readily than PLV and alga correlations. We attribute this to the likely rapid proliferation of the *Nucleocytoviricota* groups after infection (*e*.*g*., *Chlorella* virocells release PBCV-1 viral particles three hours post infection (Van Etten et al., 1983)). In the case of PLVs, although relatively little is known, their dual lifestyle (Krupovic and Koonin, 2015; Yutin et al., 2015) of insertion, and later proliferation may require specific cues for a virocell to develop and release viral particles effectively uncoupling infection and release of viruses acutely, thereby confounding the interpretation of a correlation analysis. Although *Lavidaviridae* could be uncoupled in the same way due to viral integration capabilities, in this case a known trigger for viral particle production is present (*i*.*e*., co-infection with *Nucleocytoviricota*) and conclusively we expect all members of this tripartite infection to co-exist when *Lavidaviridae* are detected in the HRAP. In this scenario, detection of the *Lavidaviridae* indicates viral particles being produced and this would be in response to cellular hosts with a *Nucleocytoviricota* infection taking place (*i*.*e*., a virocell is formed). With ecology in mind, we expect to be able to produce some valid indication of both *Nucleocytoviricota* and *Lavidaviridae* alga hosts through correlation analyses, however PLV hosts indication are less straight forward. Additionally, some interactions could be occurring between PLVs and *Nucleocytoviricota*, but these were not explored given the lack of ecological information on these putative PLVs.

Using our full qPCR dataset (**Figures 3** and **4**) and Chlorophyta metabarcoding results (**Figure 5**) we can further hypothesize virus-host interactions by using our correlation and clustering work as an initial indication of potential hosts. Overall viruses within hierarchically clustered groupings exhibited similar patterns of presence and absence based on qPCR results in both *Nucleocytoviricota* and *Lavidaviridae* (**Figure 3**), and PLV analyses (**Figure 4**). Notably, the *Nucleocytoviricota* and *Lavidaviridae* grouping (**Figure S6 b**) reflect *Picochlorum* ASVs quite closely (**Figure 5 F**), strengthening our hypothesis that members from this grouping could be infecting members of genus *Picochlorum*. Although overlap does occur between the single member of *Mimiviridae* in group d (**Figure 3**) and correlated algae from genera *Dictyosphaerium*, and *Mantoniella* (**Figure S7 c,d**), no obvious clarification is made by the full dataset and qPCR implied abundances are quite low. Finally, the single *Mimiviridae* in group e (**Figure S7**) has further evidence for infecting the “unassigned” *Chlorella* species (**Figure 5 F**), as previously indicated by our correlation analyses, given both of their nearly complete disappearance after May 2018.

**Figure 3.**
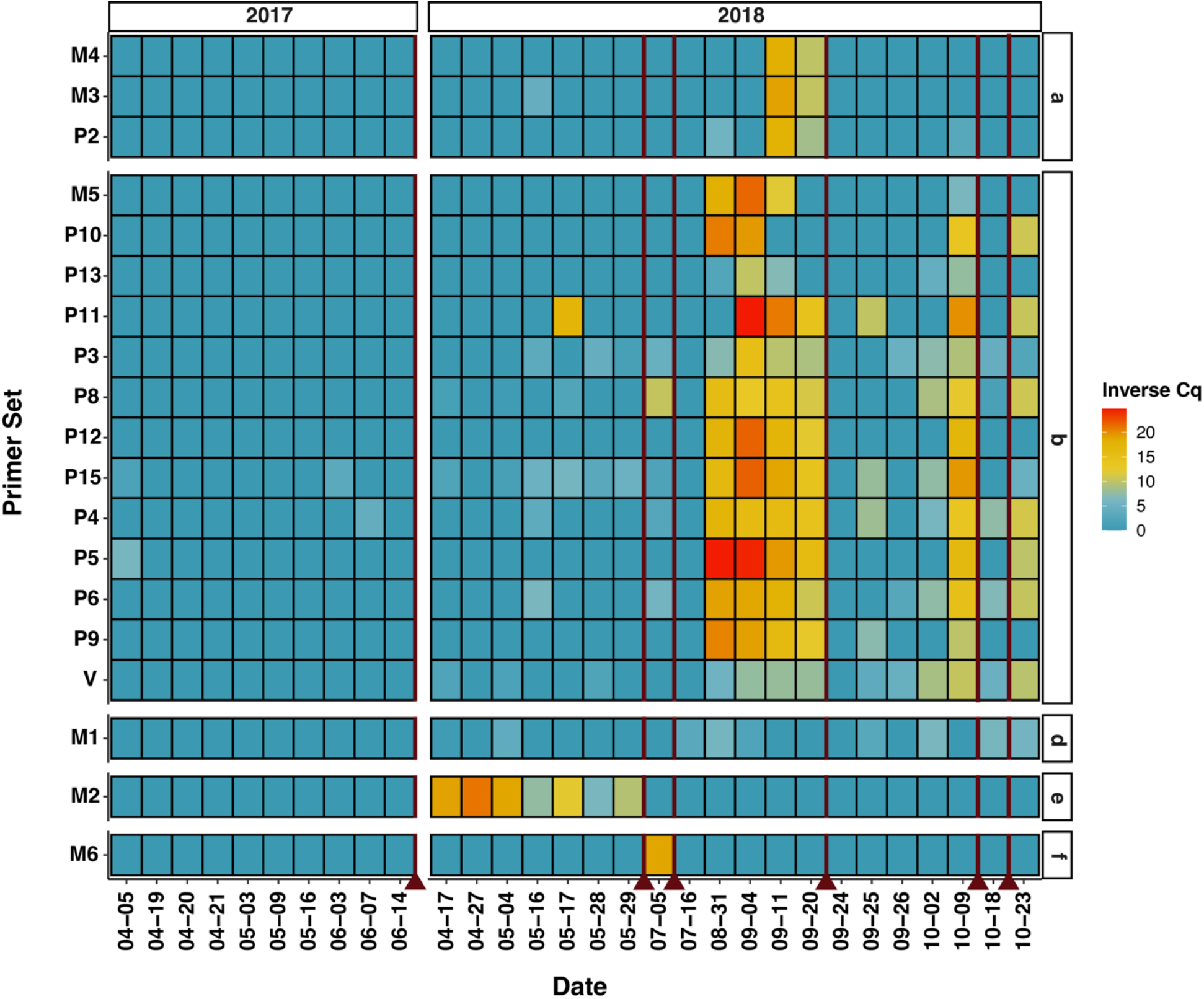
Putative Nucleocytoviricota and virophage tracking by qPCR throughout 2017 and 2018 samples. Where V, P, and M indicate potential virophage (family *Lavidaviridae*), family *Phycodnaviridae*, and family *Mimiviridae* respectively. An inverse Cq is calculated by number of cycles (total 45) minus the mean Cq value across technical triplicate reactions for each instance (*e*.*g*., viral target and sample data). Redlines indicate HRAP crash dates. Note, group c is not present given that no viruses were included during hierarchical clustering.

**Figure 4.**
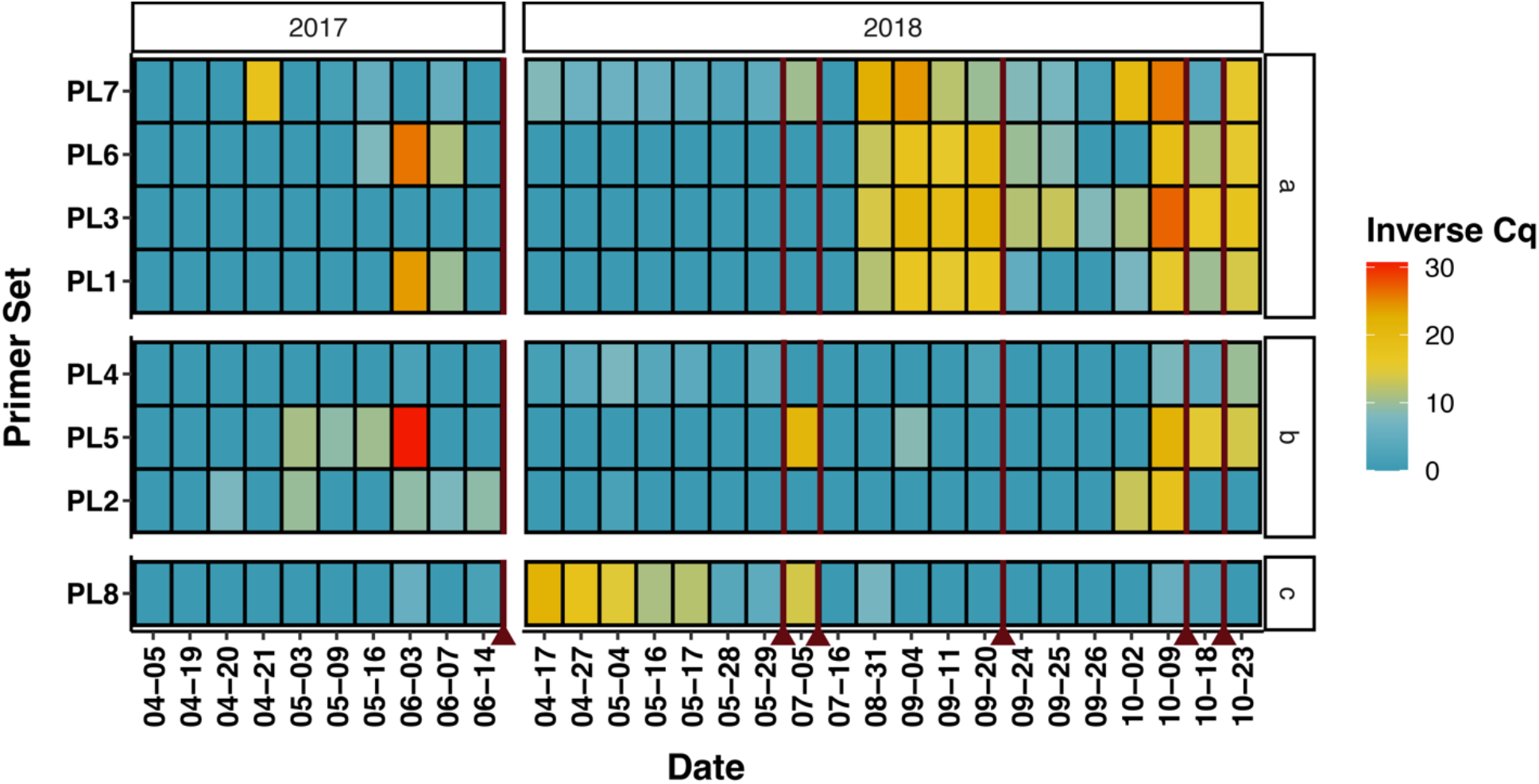
Putative polinton-like virus (PLV) tracking by qPCR throughout 2017 and 2018 samples. Where PLV indicates potential polinton-like viruses. An inverse Cq is calculated by number of cycles (total 45) minus the mean Cq value across technical triplicate reactions for each instance (*e*.*g*., viral target and sample data). Redlines indicate basin crash dates.

**Figure 5.**
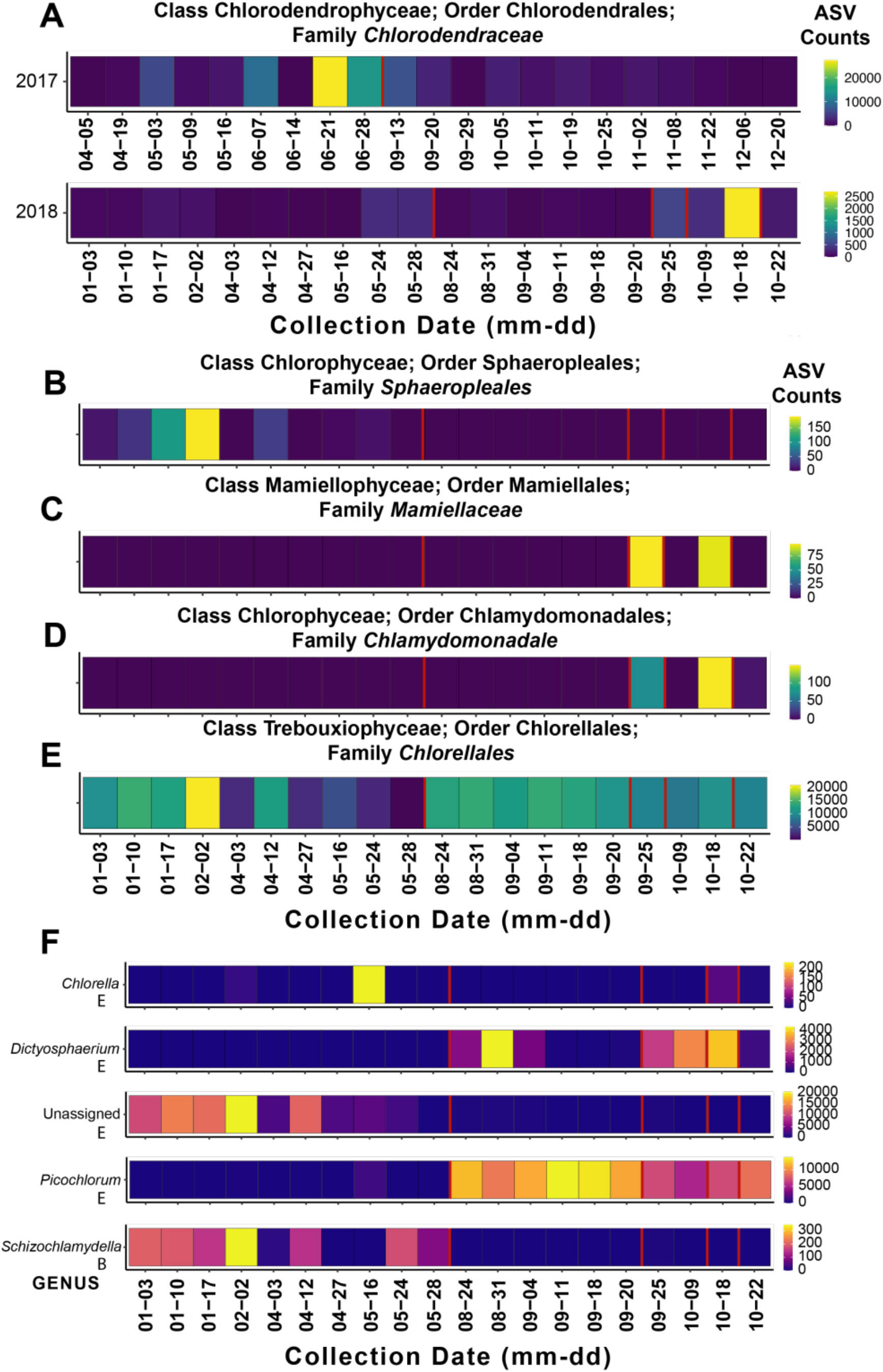
Presence and relative amounts (based on amplicon sequence variants (ASVs) of 18S data) of potential microalgae hosts using 2018 data (with the exception of 2017 being included in **A**) collected by water samples of the basin system. **(A-E)** The most abundant potential microalgal hosts at the Family level are shown, and **(F)** Genus level for family *Chlorellales* and family *Sphaeropleales* (as indicated by E or B). ASV counts are indicated by colour legends. Instances where a new culture run is made, after a die-off, are indicated by a red line.

Interestingly, the PLVs most closely related to a known PLV infecting *Tetraselmis striata* (**Figure S3;** PLV 2, 5, and 7) do not correlate closely with *Tetraselmis* spp. in the HRAP based on our PCA analysis (**Figure S6 B**), however two out of three significantly cluster together (**Figure S8 b**). We cannot confidently conclude that none of the tracked PLVs infect *Tetraselmis* spp. not only because the theoretical decoupling of PLVs and hosts discussed above, but also because an important peak of *Tetraselmis* spp. ASVs occurs during a period where qPCR data is unavailable (mid-June to autumn 2017; **Table S3**). This peak would likely be an important part of the data to assess the specific relationship (if any) between PLV 2, 5, and 7 and *Tetraselmis* spp. Phylogenetically closely related PLVs could be infecting completely different organisms, therefore the relationship of PLVs 2, 5, 7, to TsV-N1 (and each other) does not completely confirm infection of the same species. Overall, correlating PLVs and hosts by these methods are made difficult by their ecology (and our minimal understanding of PLVs at present). No other classifications (by ASVs) outside of Chlorophyta correlate significantly (including hierarchical clustering analysis) with these PLVs (data not included), despite the discovery of PLVs infecting several groups outside of Chlorophyta (Bellas and Sommaruga, 2021). However, alternative hosts (*i*.*e*., not members of Chlorophyta) cannot be ruled out by these *in silico* analyses alone. An important point is that viruses found in different taxonomic groups can also be infecting the same organism (Bellas and Sommaruga, 2021), consequently it is difficult to infer a host of these tracked PLVs. Regardless of the true host, this HRAP appears to be an interesting and significant reservoir of PLVs, some quite likely to be infecting microalgae.

Given the similarity of *Picochlorum* (**Figure 5 F**) and group c virus (**Figure 3**) dynamics (*i*.*e*., presence and abundance) with visual inspection and the results of our correlation analyses, the possibility of a previously undocumented infection of *Nucleocytoviricota* and a tripartite infection with a *Lavidaviridae* to a *Picochlorum* required further investigation. Analysis of eukaryotic genomes has revealed that endogenous *Nucleocytoviricota* are common in some lineages, particularly in green algae, and thus play an important role in host genome evolution (Blanc et al., 2015; Gallot-Lavallée and Blanc, 2017; Moniruzzaman et al., 2020). These viral insertions provide evidence of past interaction between the original virus and the organism that captured its DNA. Available *Picochlorum* genome assemblies were downloaded from NCBI GenBank to check for *Nucleocytovirivota* integrated *Nucleocytoviricota* genes/viral elements (GenBank accessions GCA011316045, GCA010909725, GCA00281821, GCA009650465, GCA_000876415). Several regions with *Nucleocytoviricota* signatures were recovered among these genomes (median size of ∼15 kb; **Table S4**) using the dedicated ViralRecall program (Aylward and Moniruzzaman, 2021), interestingly a *Picochlorum* genome recovered from the Mediterranean Sea (Costa Vermeille) (Krasovec et al., 2018) possessed a relatively large region (∼94 kb) with both superfamily II helicase and D5 primase-helicase markers detected. These results suggest that *Nucleocytoviricota* do or have been infected with *Picochlorum* in the past (this particular sample was taken in June 2011), and specifically in the geographical area where the HRAP was situated.

### Diversity of Phycodnaviridae may reflect diversity of the host

Of substantial interest in this study is the scale of *Phycodnaviridae* (phycodnaviruses) uncovered in our MVs (**Figure 6**). *Mimiviridae* representatives also appear, although in less abundance. It is known that *Phycodnaviridae* genomes of genera *Prasinovirus* and *Chlorovirus* contain multiple copies of the MCP gene (*i*.*e*., paralogs) (Clerissi et al., 2014), therefore the sheer number of MCP hits in this phylogeny may be at least partially explained by intragenome duplication. Nonetheless, as a core gene of *Nucleocytoviricota* (Yutin et al., 2009), putative MCPs are a vital finding in the HRAP because they signal true viral hits. With respect to DNA polymerase (PolB), a relatively robust single-copy gene for phylogenetic reconstruction of *Nucleocytoviricota* (Chen and Suttle, 1996; Clerissi et al., 2014), there are still a considerable number of putative *Phycodnaviridae* (19 total; **Figure S9**). Finally, a considerable number of ATPase sequences (another marker of *Nucleocytoviricota*) with *Phycodnaviridae* hits are also recovered from the MVs (18 total; **Figure S10**). Our results appear to suggest that in an environment such as an HRAP, where a dense culture of microalgae simulates an algal bloom, a viral “bloom” reflecting the algal abundance can occur. In a similar fashion to the RNA quasispecies (Andino and Domingo, 2015), the HRAP *Phycodnaviridae* form a repertoire of closely related genomes. RNA quasispecies are said to involve a high copy number of genome variants arising from the high mutation rates of RNA viruses (Duffy, 2018). This phenomenon produces so-called “mutant swarms” that foster unique dynamics between viruses where variants are acting “within replicate complexes, within cells, or outside cells” (Domingo et al., 2012; Domingo and Perales, 2019) in competition amongst themselves. The results of this, to the host(s)’ detriment and the virus population’s benefit, is the production of a “genome repertoire” (Domingo et al., 2020) in advance of a host’s response, or other factors that could affect the host-virus dynamics of a system in the future. However, instead of multiple viral, quasi-identical strain-like species of the RNA quasispecies, HRAP phycodnaviruses exhibited substantial genomic sequence divergence given dotplot alignments of contigs containing a single-copy DNAP gene (**Figure S11**). RNA viruses are capable of evolving as quasispecies because their error-prone (*i*.*e*., low fidelity) RNA polymerase, resulting in a mutational rate up to 1 million times that of their hosts (Duffy, 2018). *Nucleocytoviricota* do not feature the same high rate of mutation, largely attributed to high fidelity PolB repair machinery (Fischer et al., 2014; Redrejo-Rodríguez and Salas, 2014). Thus, it seems likely that the phycodnaviral lineages had begun to diverge well before they entered in the culture basin and on a larger time scale than between members of a typical RNA quasispecies. Assuming that the HRAP phycodnaviruses infect the same species they would have to compete for access to the host. Under competitive conditions, the maintenance of a large diversity of viruses is not the most likely scenario, since ecological theory rather predicts the hegemony of the most fit virus. A possibly more likely scenario would be that the different viral lineages have evolved different strain specificity within the same host species. Under the very favourable condition of a HRAP culture, multiple algal strains could co-exist durably and support replication of a diversity of related viruses. Alternatively, this diversity could reflect the diversity of closely related species present within one alga genus (*i*.*e*., *Picochlorum*).

**Figure 6.**
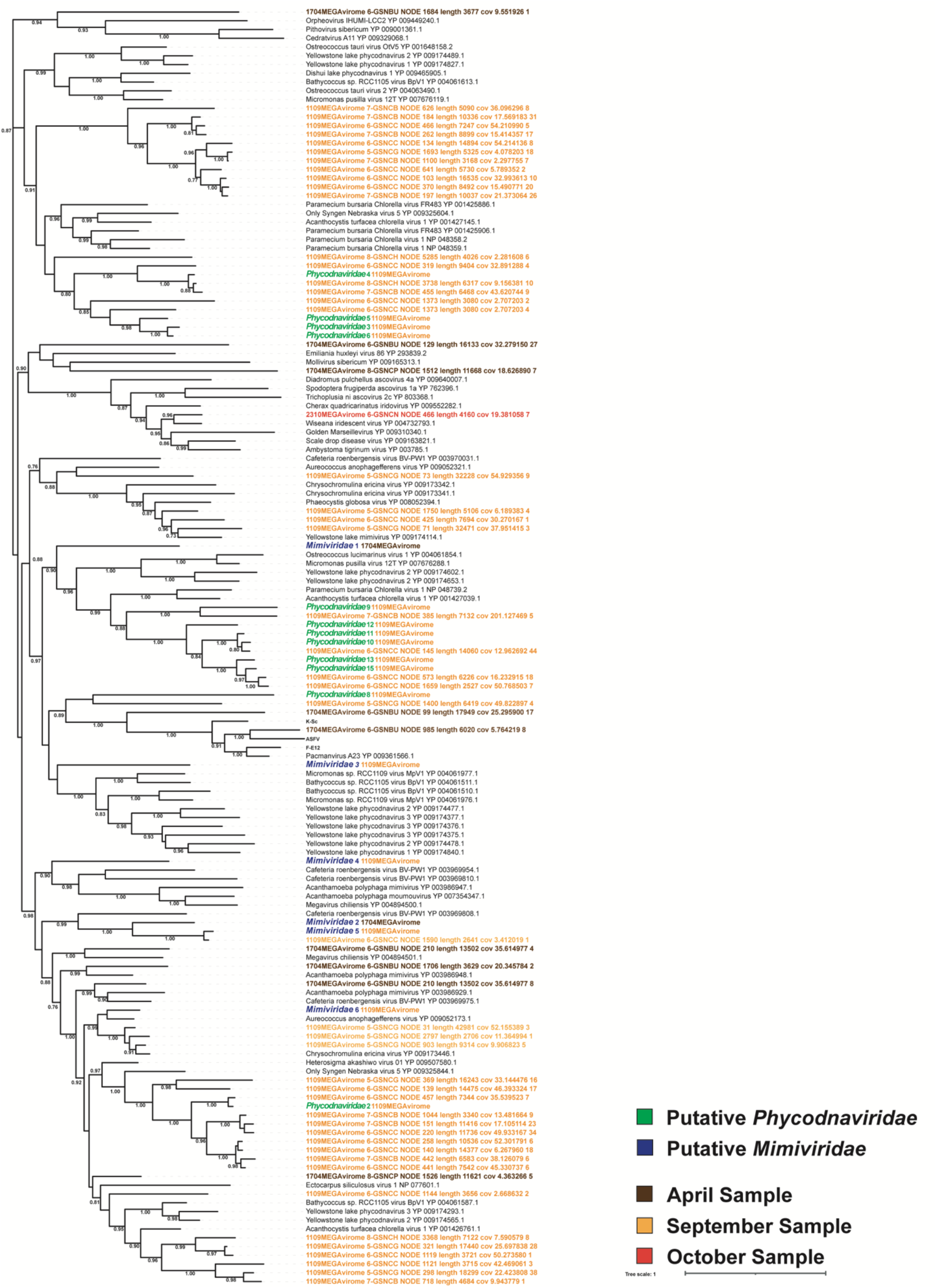
Phylogenetic relationship of *Nucleocytoviricota* found within the study (denoted as “unclassified”), and sequences obtained from NCBI GenBank (accession numbers as indicated) based on major capsid protein (MCP) amino acid sequence. Groups are assigned based on clades of closely related (by sequence similarity) novel *Nucleocytoviricota* for further study. Bootstrap support is indicated and based on 1000 replicates.

Within RNA quasispecies, diverse genotypes are permissible in order to handle the diversity of strains occurring within their host target species, such as the bloom species *Heterosigma akashiwo*, which has a wide distribution (Lawrence and Suttle, 2004). An array of virus strains is kept as a repertoire of infectious agents to handle the dynamics of their host strains. It may not be the case that new algal strains are developing, but rather a dance between dominant strains, and strains that are simply maintained, which changes over time or space. Although the exact mechanism is not the same between the HRAP phycodnaviruses and the RNA viruses, the pressure put on the viruses to evolve (*i*.*e*., the presence and maintenance of specific traits of a species pangenome) and the evolutionary response, or “solution”, are similar. The said pressure being the dynamics and repertoire of microalgae strains and closely related species (in our case members of family *Chlorellales –* potentially genus *Picochlorum*), and the solution being having a complementary genome repertoire of related viruses at hand; in other words a genome repertoire to handle host diversity similar to RNA quasispecies evolutionary “behaviour”. In a study of *Prasinoviruses* found in *Ostreococcus* spp. (Yau et al., 2018), authors investigated a mechanism of dynamics by which *Nucleocytoviricota* and their hosts interactions centred around the host species (a microalgae species or strain) being resistant or susceptible to virus infection. Simplified, it was shown that in different environments, and in presence or absence of viruses, different microalgal life history strategies involving virus-susceptible and virus-resistant microalgae would be favoured or disfavoured. We thereby suggest that these phycodnaviruses forming this level of diversity in association with a dense bloom could be defined as a “quasigenus”, akin to the RNA quasispecies. This quasigenus consist of a group of related, but individual distinct, *Phycodnaviridae* that are capable of maintenance during a dense bloom of their host alga. Alga strains within one alga species can be virus susceptible or resistant (*e*.*g*., with the system of *Aureococcus anophagefferens* and AaV (Gobler et al., 2007; Brown and Bidle, 2014; Gann et al., 2022)), thereby a dense culture with different strains could support a group of closely related viruses (the “quasigenus”) that are capable of infecting some (one or more) strains of an alga species and not others. It will be interesting to determine if this “quasigenus” phenomenon also happens so prominently during algal blooms in natural environments or if it meets only in the very favourable conditions of the HRAP. Although the concept of several different viruses infecting one species is not new, it is specifically intriguing that the maintenance of several phycodnaviruses could reflect the persistence of algal strains within possibly one host species.

### Viruses may be a factor of mass mortality and bloom control within the HRAP

Several culture crashes occurred in the HRAP throughout 2017 and 2018, some were during periods where our dataset is not at a high enough resolution to make implications about whether viruses were a significant factor or not. For example, *Mimiviridae* 1 (**Figure 3 e**) and PLV 8 (**Figure 4 c**) are the only viruses tracked that seemingly persists before the July 2018 crash (excluding 2017 data), although not many samples were collected near the July 2018 crash. The resolution around the three crashes occurring in Autumn 2018 are quite well defined overall, and consequently the abundance of viruses (**Figure 3** b, **Figure 4 a**) appear to coincide well with these crashes, thereby we hypothesize that the group of mostly phycodnaviruses are contributing to the crash of primarily *Picochlorum*. This is not unusual as viruses are known to participate in bloom termination (Gastrich et al., 2004; Lawrence and Suttle, 2004; Brussaard et al., 2005). In context of the co-occurring PLVs, it is possible they are simply present and absent based on the presence and the removal of their potential hosts (*e*.*g*., induced by other predators, parasites or fluctuating seasonal conditions) and the overall crash and re-initiation of the system, but more significantly we do not know the effect of these PLVs on their hosts and if PLVs can contribute to or cause bloom termination in marine environments (*e*.*g*., if these PLVs lyse their hosts versus behave like *Lavidaviridae* with a tripartite infection system). Broadly speaking, the HRAP may not completely reflect the trajectory of Mediterranean sourced water used to initiate the HRAP culture, given that some algae only appear around times of culture restarts and then disappear after (family *Mamiellaceae* and *Chlamydomonadale*; **Figure 5**), however for the microalgae that persist and the viruses occurring alongside there appears to be an algal bloom like maintenance and resulting virus bloom termination.

## Conclusion

Overall, this study demonstrates the incredible diversity and dynamics of DNA viruses within an HRAP. Ultimately, we suggest that the HRAP environment can mimic a marine microalgae bloom. This resulting diversity of putative *Phycodnaviridae* contains a repertoire of viruses available to infect, at least one genera (and potentially the strains of one or more species) of microalgae, giving way to the concept “quasisgenus” of viruses. Although the exact progression to reach this result are different than that of RNA quasispecies, we cannot deny similarities in the end results of this potential virus-host dynamic. Ultimately, it appears that viruses (specifically those involved in a “quasisgenus” behaviour) could be meaningfully contributing to the termination of an algal bloom in the HRAP system.

## Supporting information

Table_S3_ASV_counts

Table_S1_Assembly_statistics

Table_S4_ViralRecall_Picochlorum_viral_insertion_data

Table_S2_Functional_annotations

## Abbreviations

(HRAP): High-rate algal pond
PLVs: Polinton-like viruses
FACS: fluorescence activated cell sorting
UV: ultraviromes
MV: Megaviromes
(ORF): open reading frame
MCP: major capsid protein
PolB: DNA polymerase B
PCA: principal component analyses
SIMPROF: similarity profile analysis
AMG: auxiliary metabolic genes
RM: restriction modificatino

## Acknowledgments

The authors thank the members of the VASCO2 consortium and the company COLDEP for their support and permission to access the cultures. We also acknowledges support by the Institut Français de Recherche pour l’Exploitation de la Mer (IFREMER) and work done by Angélique Gobet. Including the Ifremer Palavas-les-Flots raceway facility and GeT-PLaGe platform (Genotoul). We also wish to specifically acknowledge our late co-author, Dr. Christelle Desnues, who brought compassion, excellence, and inspiration to research during her life.

## Author Contributions

Conceptualization, C.D., G.B., E.E.C., and S.M.-B.; Methodology, C.D., G.B., S.M.-B., E.E.C.; Software, E.E.C., and G.B.; Validation, formal analyses, and investigation, G.B., C.D. and E.E.C.; Resources, C.D. and G.B.; Data Curation, E.E.C. and G.B.; Writing—Original Draft Preparation, E.E.C.; Writing—Review & Editing, G.B., C.D., S.M.-B., T.M.P, and E.E.C.; Visualization, E.E.C.; Supervision, C.D., G.B., and S.M.-B.; Project Administration, C.D., and G.B.; Funding Acquisition, G.B., and C.D. All authors have read and agreed to the published version of the manuscript.

## Funding

E.E.C, G.B, and C.D received funding from the European Union’s Horizon 2020 research and innovation programme under the Marie Skłodowska-Curie grant agreement No713750, with the financial support of the Regional Council of Provence-Alpes-Côte d’Azur and with the financial support of the A*MIDEX (n° ANR-11-IDEX-0001-02), funded by the Investissements d’Avenir project funded by the French Government, managed by the French National Research Agency (ANR). The Phycovir project leading to this publication has received funding from Excellence Initiative of Aix-Marseille University—A*MIDEX, a French “Investissements d’Avenir” program. Part of the sequencing was conducted by the U.S. Department of Energy Joint Genome Institute under CSP Proposal: 10.46936/10.25585/60000928, a DOE Office of Science User Facility (https://ror.org/04xm1d337), is supported by the Office of Science of the U.S. Department of Energy operated under Contract No. DE-AC02-05CH11231.

## Data Availability

Data are available through and hosted by the NCBI SRA portal under the BioProject I PRJNA751746 (MiSeq ultraviromes), and by the JGI Genome Portal under the JGI Proposal Id: 504991 (HiSeq megaviromes).

## Supplementary Materials

Table S1: Assembly statistics, Table S2: functional annotation of viral contigs, Table S3: ASV counts, Table S4: information on predicted viral regions in *Picochlorum* genomes. Figure S1: Fluorescent activated cell sorting of megaviral populations, Figure S2: Broad taxonomic classification of contigs for all sequencing dataset, Figure S3: Putative polinton-like viruses (PLVs) recovered from HRAP metagenomes, Figure S4: Putative *Lavidaviridae* assembled from HRAP assemblies, Figure S5: Principle component analyses, Figure S6: Hierarchical clustering of potential hosts and viruses. Figure S7. Hierarchical clustering of potential hosts and polinton-like viruses (PL). Figure S8: Putative *Nucleocytoviricota* phylogeny based on the DNA polymerase B gene, Figure S9. Putative Nucleocytoviricota phylogeny based on the ATPase gene. Figure S10. Dotplot alignments of *Phycodnaviridae* contigs containing a PolB gene.

## Supplemental Figures

**Figure S1.**
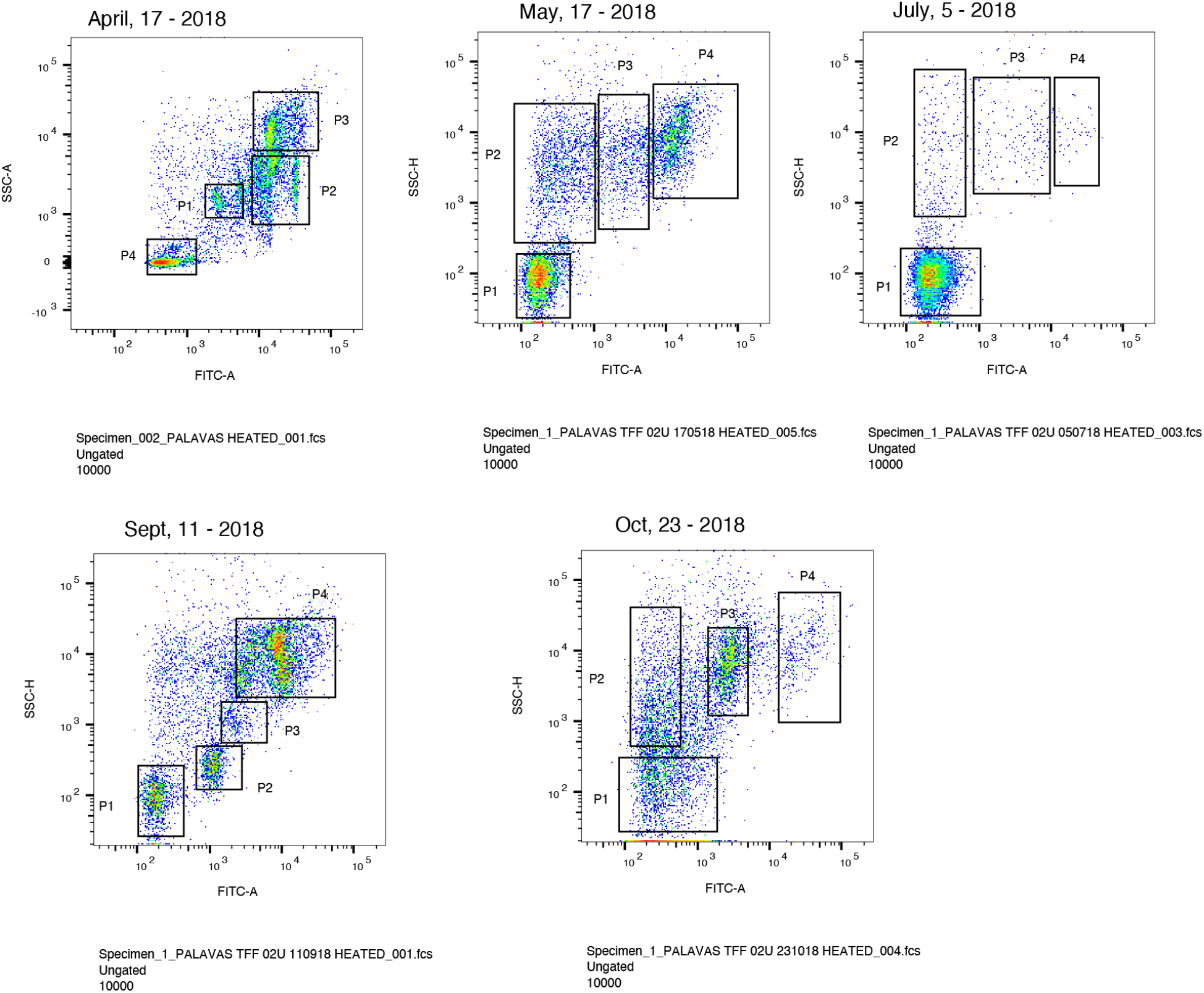
Fluorescent activated cell sorting by side scatter (SSC) and FITC (fluorescein isothiocyanate; excitation and emission spectrum peak wavelengths of ∼495 nm and ∼519 nm). Sample dates are displayed above each sample, and populations (“P”) are indicated. The sized fraction for the sample water is between 0.2 μm –1.2 μm (labelled with SYBR-Green).

**S2.**
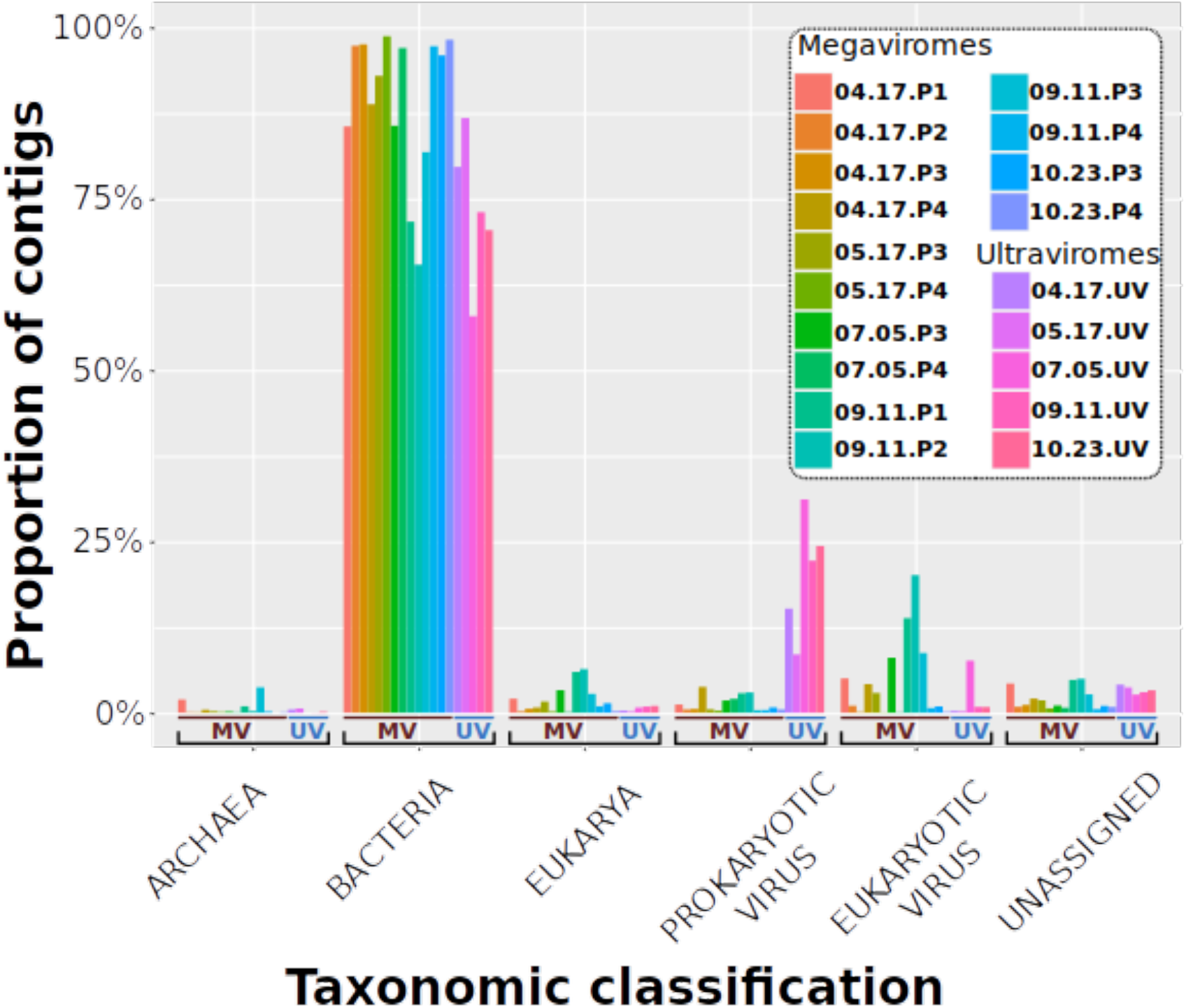
Broad taxonomic classification of contigs above 2 kb for all megaviromes sequenced and a combined ultravirome sequencing dataset.

**Figure S3.**
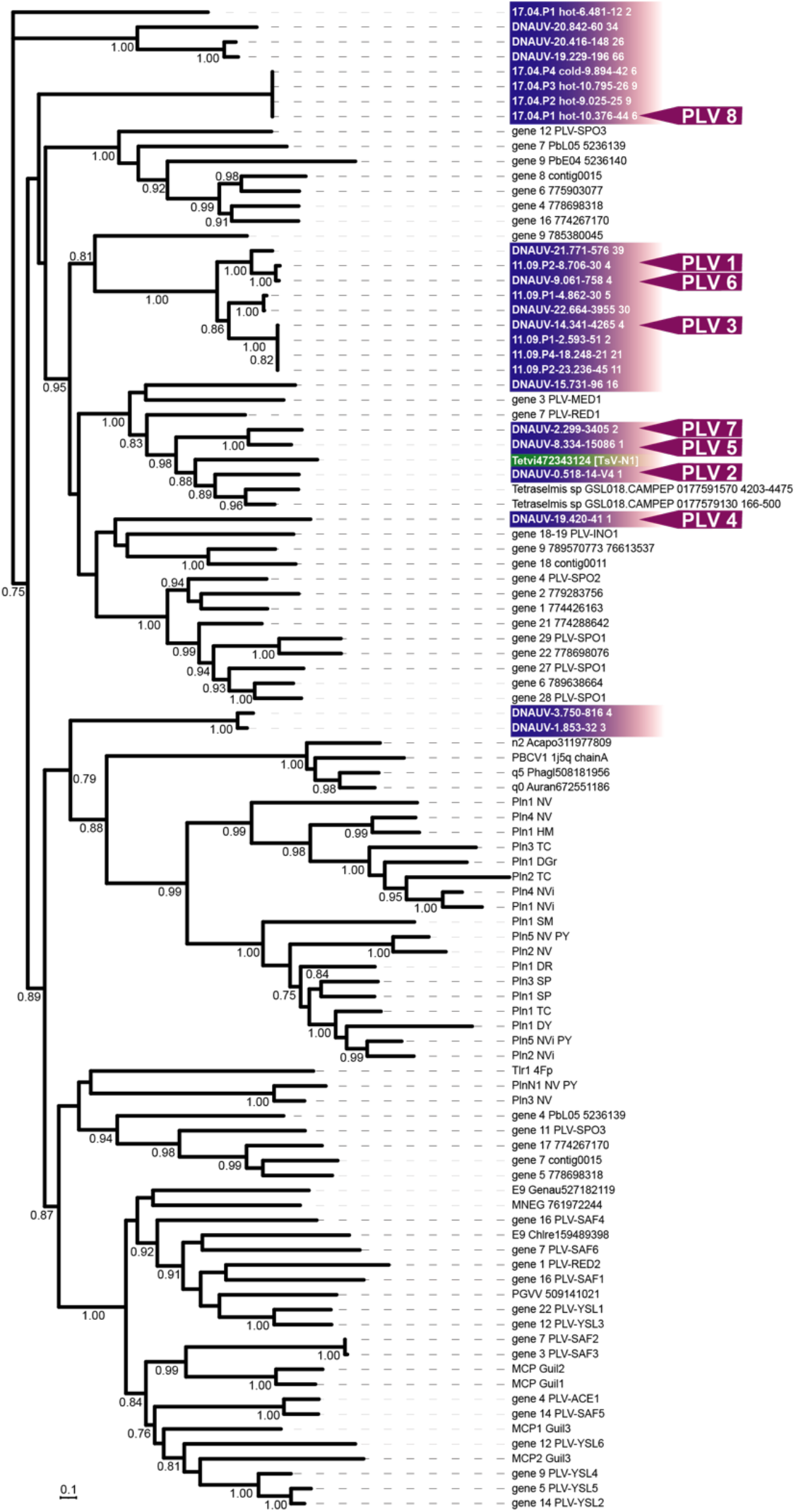
Putative polinton-like viruses (PLVs) recovered from HRAP metagenomes (indicated in red). The PLV TsV-N1 is indicated and the green line covers the PLVs that are clustering with it phylogenetically. The phylogeny based on the MCP gene. PLVs used in qPCR tracking are indicated in blue. Pol 8 is excluded due to the sequence dissimilarity relative to the rest of the tracked PLVs. Bootstrap values above 0.70 are displayed. All MCP (excluding those in red) were extracted from public databases (see Chapter 3 for more detailed methods).

**Figure S4.**
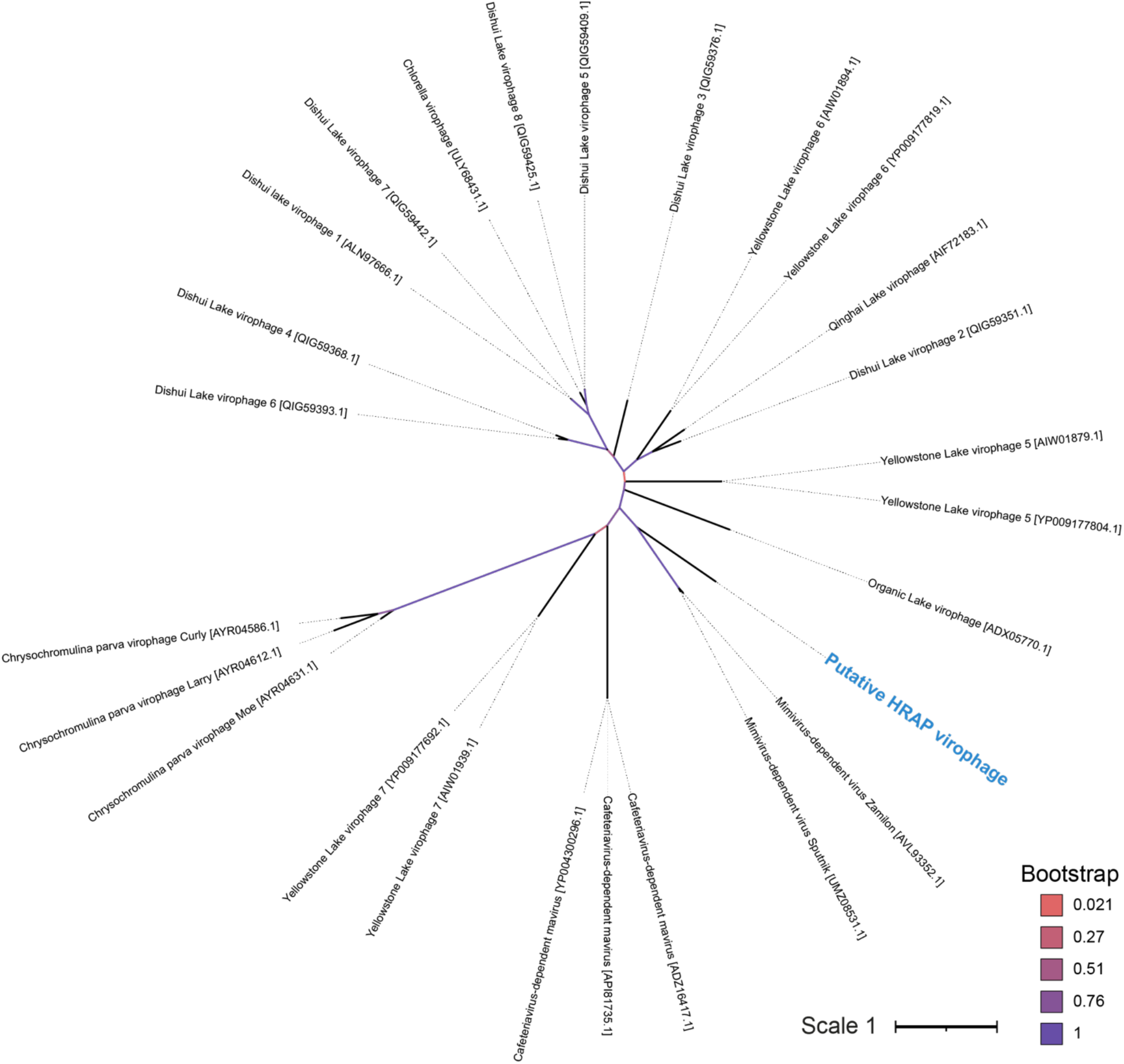
Putative *Lavidaviridae* (in blue) assembled from HRAP assemblies. Phylogeny is based on major capsid protein sequenced retrieved from NCBI GenBank. Bootstrap probabilities (1000 replicates) are displayed.

**Figure S5.**
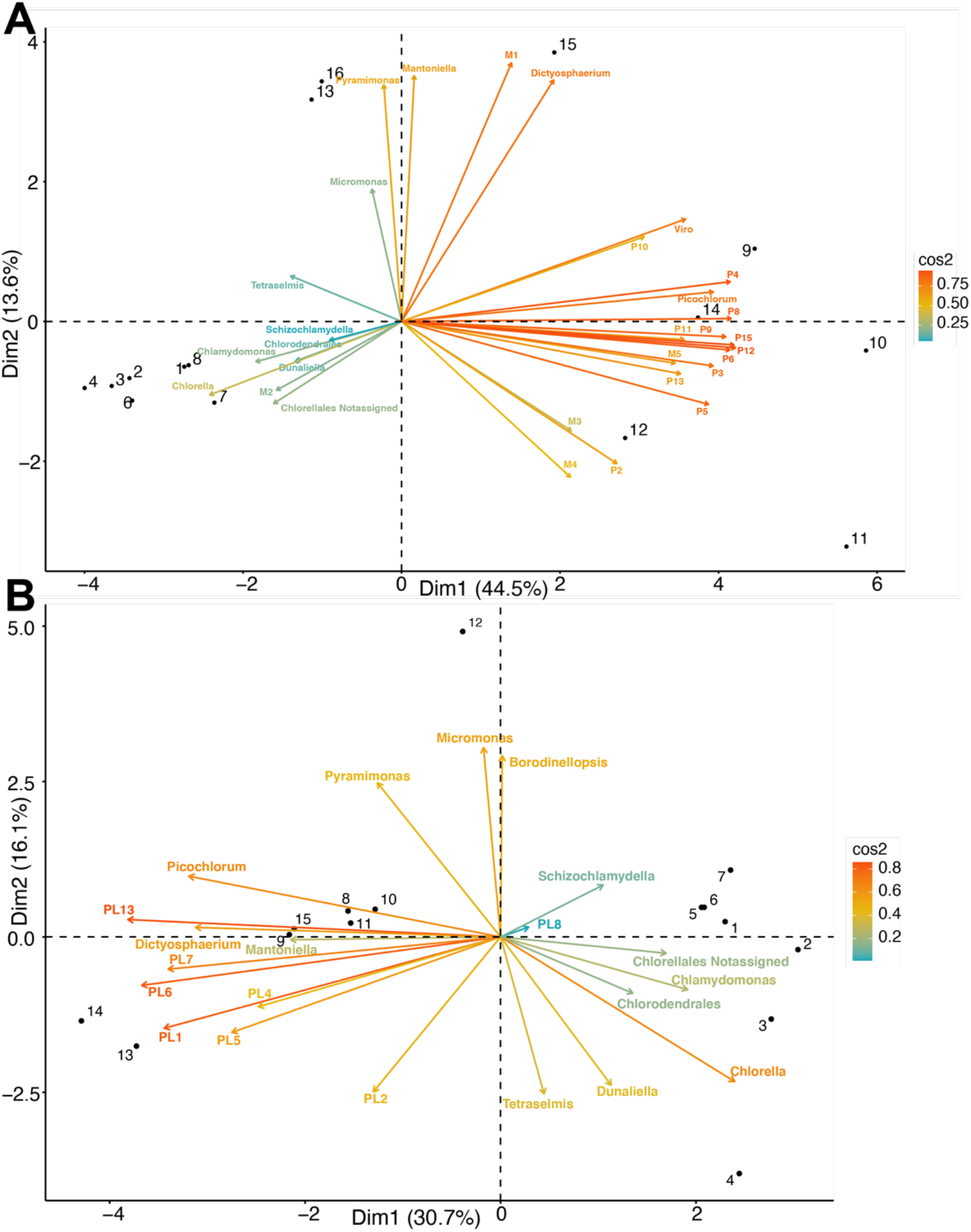
Principle component analyses using combined and normalised qPCR data and 18S ASVs (i.e. metabarcoding) on tracked putative viruses and potential alga (Chlorophyta) hosts for **(A)** *Mimiviridae* (M), *Phycodnaviridae* (P), and virophage (Viro), and **(B)** polinton-like viruses (PL). Cos2 reports the strength of the principle component for the observations (i.e. virus or potential hosts), where a higher value depicts a stronger relationship between them or a “good representation”. The vector length of each observations represents the contribution they make to the ordination. Dates are represented by numbered objects, where 2017 is composed of 1–5 (April is 1 and 2, May is 3, June is 4 and 5), and 2018 is composed of 6–16 (April is 6, May is 7– 9, August is 9, September Is 10–13, and October is 14–16).

**Figure S6.**
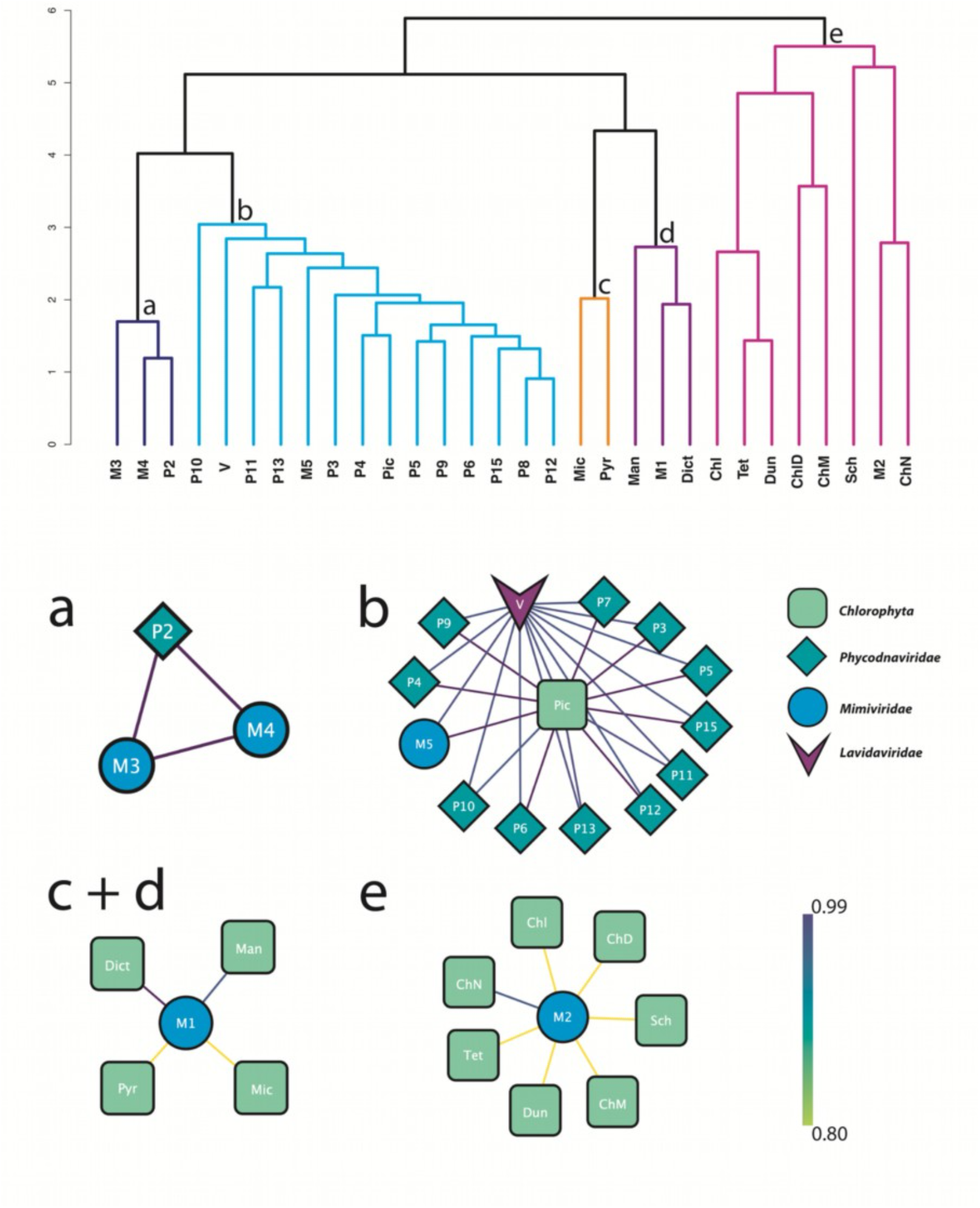
Hierarchical clustering (SIMPROF, □ = 0.05) of potential hosts (Chlorophyta) and viruses of interest; *Phycodnaviridae* (P), *Mimiviridae* (M), and virophage (V). Predicted groupings are depicted **(a–e)** using a network visualisation. Darker colours represented a higher correlation between group members, where 1 is an exact correlation and 0 is no correlation. Chlorophyta species are abbreviated as follows: Pic; *Picochlorum*, Dict; *Dictyosphaerium*, Man; *Mantoniella*, Pyr; *Pyramimonas*, Mic; *Micromonas*, Chl; *Chlorella*, ChD; *Chlorodendrales*, ChN; *Chlorella* not assigned, Sch; *Schizochlamydella*, Tet; *Tetraselmis*, Dun; *Dunaliella*, and ChM; *Chlamydomonas*.

**Figure S7.**
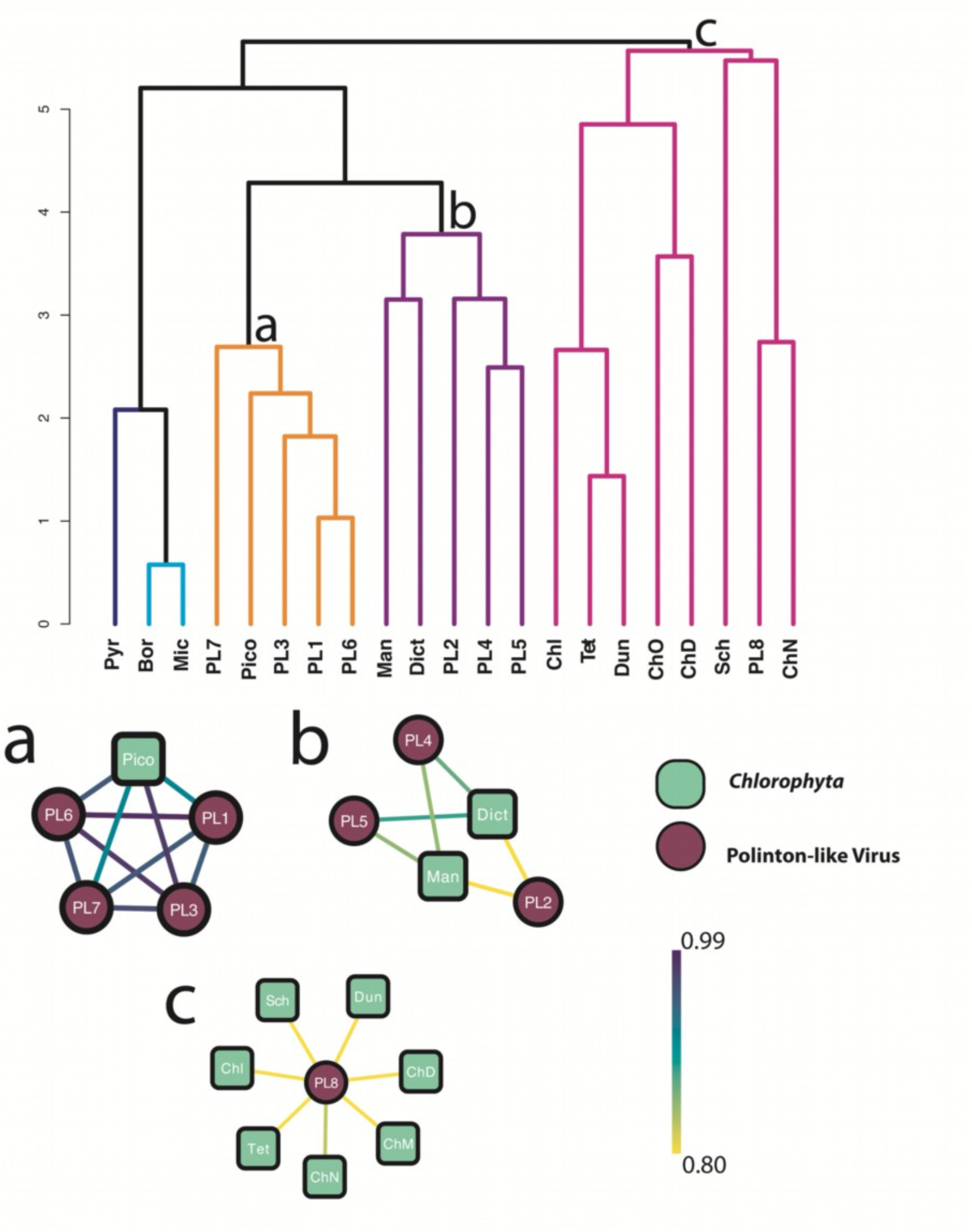
Hierarchical clustering (SIMPROF, □ = 0.05) of potential hosts (Chlorophyta) and viruses of interest; polinton-like viruses (PL). Predicted groupings are depicted **(a–e)** using a network visualisation. Darker colours represented a higher correlation between group members, where 1 is an exact correlation and 0 is no correlation. Chlorophyta species are abbreviated as follows: Pic; *Picochlorum*, Dict; *Dictyosphaerium*, Man; *Mantoniella*, Sch; *Schizochlamydella*, Dun; *Dunaliella*, Chl; *Chlorella*, ChD; *Chlorodendrales*, Tet; *Tetraselmis*, ChM; *Chlamydomonas*, and ChN; *Chlorella* not assigned.

**Figure S8.**
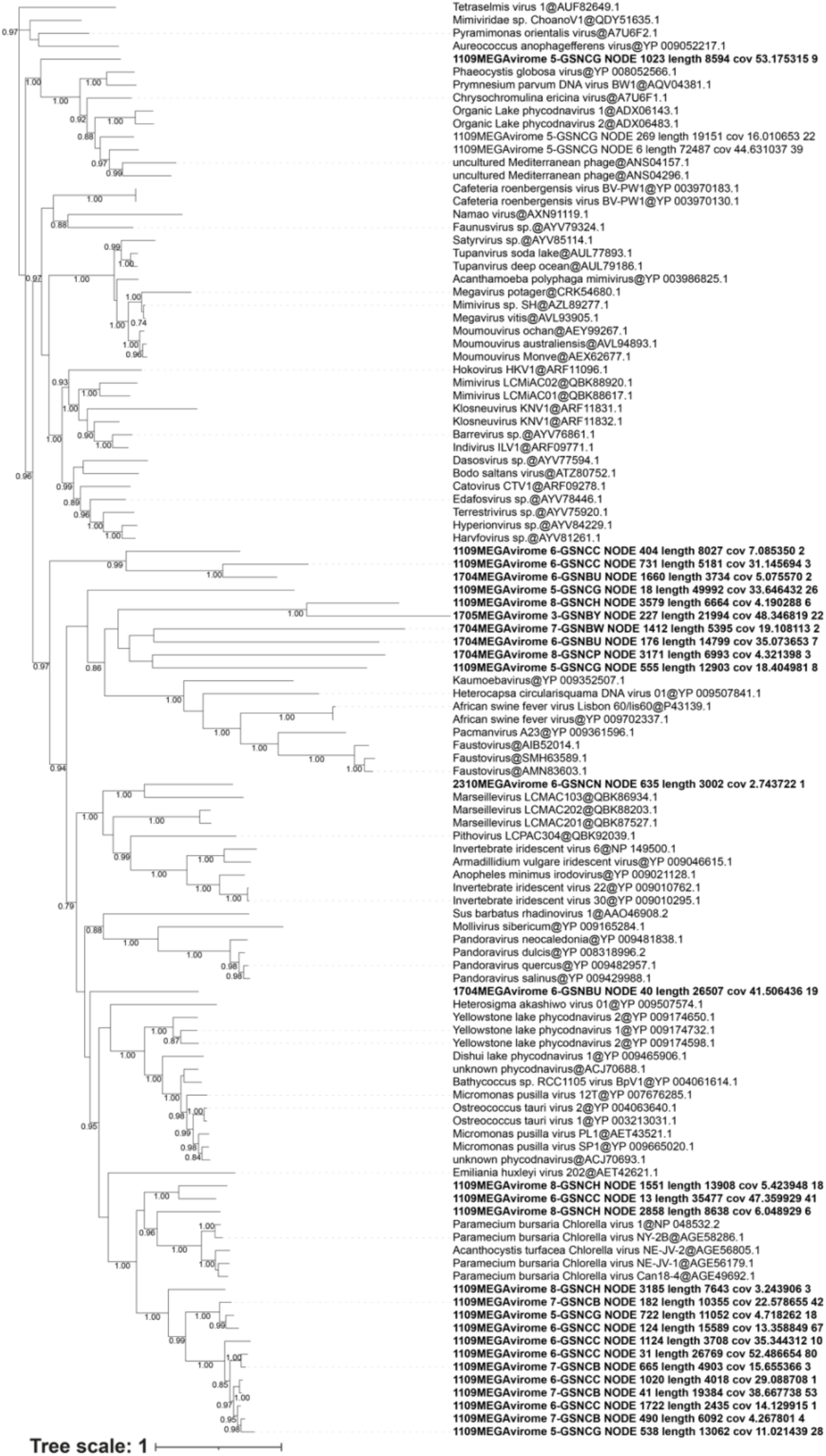
Putative *Nucleocytoviricota* phylogeny based on the DNA polymerase B gene (polB). Reference sequences were downloaded by NCBI GenBank (accession numbers as indicated) and sequenced recovered from the HRAP are included (in bold). Bootstrap values above 0.70 are displayed.

**Figure S9.**
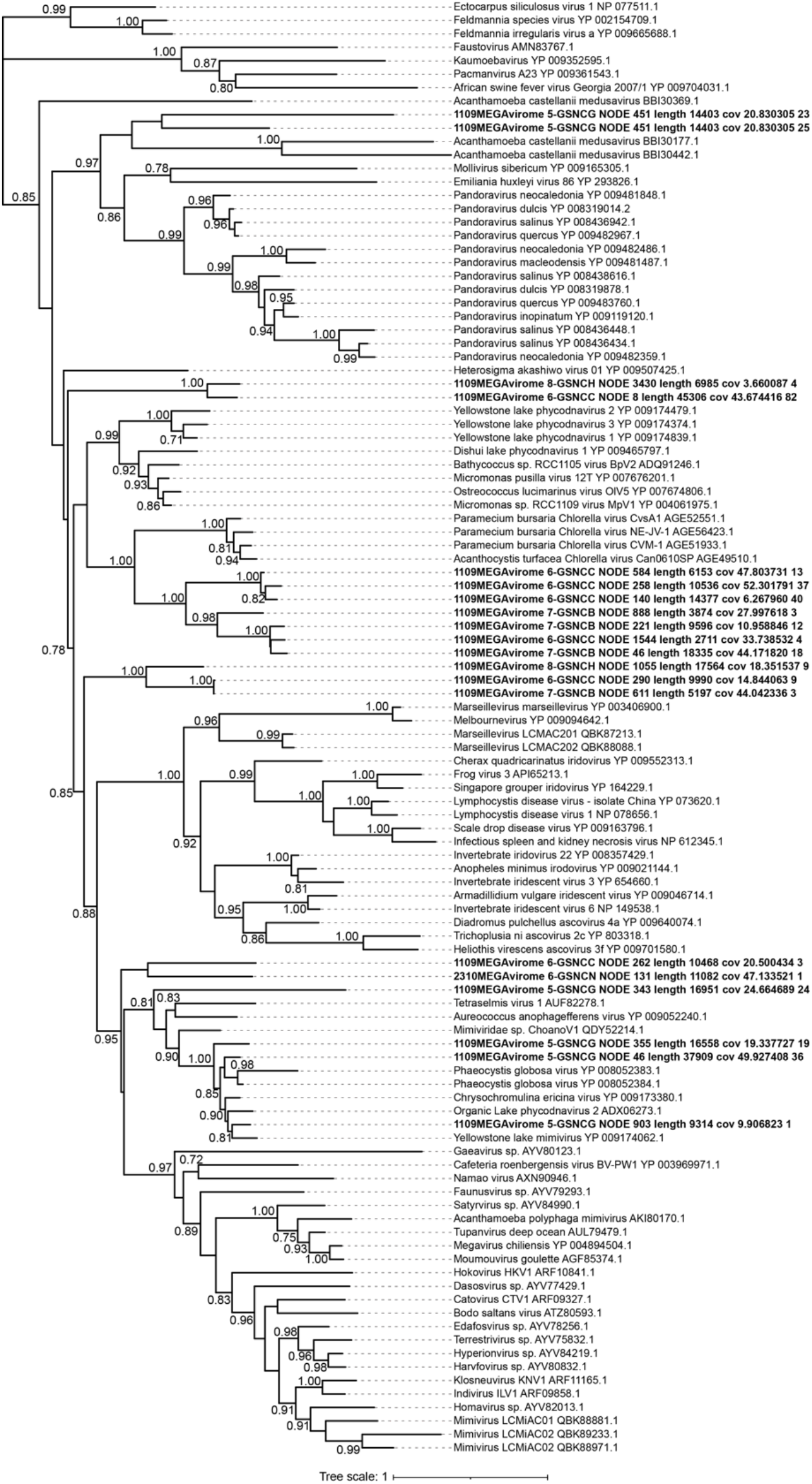
Putative *Nucleocytoviricota* phylogeny based on the ATPase gene. Reference sequences were downloaded by NCBI GenBank (accession numbers as indicated) and sequenced recovered from the HRAP are included (in bold). Bootstrap values above 0.70 are displayed.

**Figure S10.**
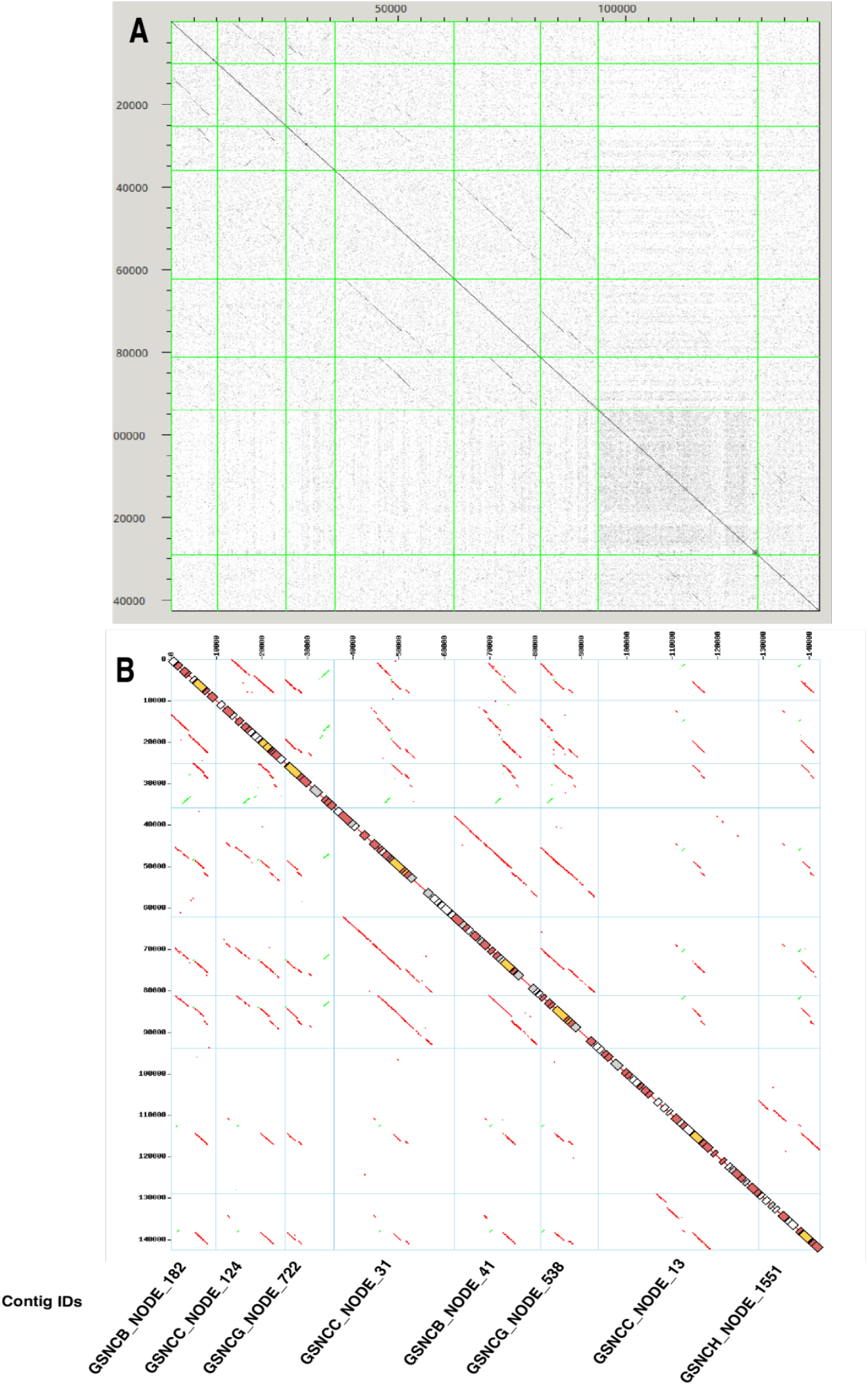
Dotplot alignments of *Phycodnaviridae* contigs containing a PolB gene. **(A)** 8 × 8 dotplot alignments of contig nucleotide sequences. **(B)** Dotplot representation of contig’s translated nucleotide sequence matches identified by TBLASTX (red for forward matches, green for reverse matches). Position of the PolB gene on individual contigs is shown by yellow rectangles on the plot diagonal. Additional ORFs with a significant match to viruses or cellular organisms in TrEMBL are shown by red and grey rectangles respectively, and ORFs > 150 codons without match are shown with open rectangles. Contig orders are identical along the x-and y-axis in **A** and **B**. Only contigs greater than 10 kb were used in the analysis. Altogether, a residual gene collinearity is generally observed between *Phycodnaviridae* contigs, with only very few inversions (green matches), however nucleotide sequence conservation is frequently interrupted outside of coding sequences indicating that intergenic sequences have diverged to the point that no significant similarity is yet detectable.

## Notes

### Competing Interest Statement

The authors have declared no competing interest.

## References

Agarkova, I. V., Dunigan, D. D., and Van Etten, J. L. (2006). Virion-Associated Restriction Endonucleases of Chloroviruses. Journal of Virology 80, 8114–8123. doi: 10.1128/JVI.00486-06.

Andino, R., and Domingo, E. (2015). Viral quasispecies. Virology 479–480, 46–51. doi: 10.1016/j.virol.2015.03.022.

Antipov, D., Raiko, M., Lapidus, A., and Pevzner, P. A. (2020). MetaviralSPAdes: assembly of viruses from metagenomic data. Bioinformatics 36, 4126–4129. doi: 10.1093/bioinformatics/btaa490.

Aylward, F. O., and Moniruzzaman, M. (2021). ViralRecall—A Flexible Command-Line Tool for the Detection of Giant Virus Signatures in ‘Omic Data. Viruses 13, 150. doi: 10.3390/v13020150.

Bellas, C. M., and Sommaruga, R. (2021). Polinton-like viruses are abundant in aquatic ecosystems. Microbiome 9, 13. doi: 10.1186/s40168-020-00956-0.

Biggs, T. E. G., Huisman, J., and Brussaard, C. P. D. (2021). Viral lysis modifies seasonal phytoplankton dynamics and carbon flow in the Southern Ocean. ISME J, 1–8. doi: 10.1038/s41396-021-01033-6.

Blanc, G., Gallot-Lavallée, L., and Maumus, F. (2015). Provirophages in the Bigelowiella genome bear testimony to past encounters with giant viruses. PNAS 112, E5318–E5326. doi: 10.1073/pnas.1506469112.

Boeckmann, B., Bairoch, A., Apweiler, R., Blatter, M.-C., Estreicher, A., Gasteiger, E., et al. (2003). The SWISS-PROT protein knowledgebase and its supplement TrEMBL in 2003. Nucleic Acids Res 31, 365–370. doi: 10.1093/nar/gkg095.

Boutet, E., Lieberherr, D., Tognolli, M., Schneider, M., and Bairoch, A. (2007). UniProtKB/Swiss-Prot. Methods Mol Biol 406, 89–112. doi: 10.1007/978-1-59745-535-0_4.

Brown, C. M., and Bidle, K. D. (2014). Attenuation of virus production at high multiplicities of infection in Aureococcus anophagefferens. Virology 466–467, 71–81. doi: 10.1016/j.virol.2014.07.023.

Brussaard, C. P. D., Kuipers, B., and Veldhuis, M. J. W. (2005). A mesocosm study of Phaeocystis globosa population dynamics: I. Regulatory role of viruses in bloom control. Harmful Algae 4, 859–874. doi: 10.1016/j.hal.2004.12.015.

Callahan, B. J., McMurdie, P. J., Rosen, M. J., Han, A. W., Johnson, A. J. A., and Holmes, S. P. (2016). DADA2: High-resolution sample inference from Illumina amplicon data. Nat Methods 13, 581–583. doi: 10.1038/nmeth.3869.

Cantalapiedra, C. P., Hernández-Plaza, A., Letunic, I., Bork, P., and Huerta-Cepas, J. (2021). eggNOG-mapper v2: Functional Annotation, Orthology Assignments, and Domain Prediction at the Metagenomic Scale. Mol Biol Evol 38, 5825–5829. doi: 10.1093/molbev/msab293.

Chase, E. E., Monteil-Bouchard, S., Gobet, A., Andrianjakarivony, F. H., Desnues, C., and Blanc, G. (2021). A High Rate Algal Pond Hosting a Dynamic Community of RNA Viruses. Viruses 13, 2163. doi: 10.3390/v13112163.

Chen, F., and Suttle, C. A. (1996). Evolutionary Relationships among Large Double-Stranded DNA Viruses That Infect Microalgae and Other Organisms as Inferred from DNA Polymerase Genes. Virology 219, 170–178. doi: 10.1006/viro.1996.0234.

Choezom, D., and Gross, J. C. (2022). Neutral sphingomyelinase 1 regulates cellular fitness at the level of ER stress and cell cycle. 2022.02.23.481585. doi: 10.1101/2022.02.23.481585.

Claverie, J.-M., and Abergel, C. (2018). Mimiviridae: An Expanding Family of Highly Diverse Large dsDNA Viruses Infecting a Wide Phylogenetic Range of Aquatic Eukaryotes. Viruses 10, 506. doi: 10.3390/v10090506.

Clerissi, C., Grimsley, N., Ogata, H., Hingamp, P., Poulain, J., and Desdevises, Y. (2014). Unveiling of the Diversity of Prasinoviruses (Phycodnaviridae) in Marine Samples by Using High-Throughput Sequencing Analyses of PCR-Amplified DNA Polymerase and Major Capsid Protein Genes. Appl. Environ. Microbiol. 80, 3150–3160. doi: 10.1128/AEM.00123-14.

Coy, S. R., Gann, E. R., Papoulis, S. E., Holder, M. E., Ajami, N. J., Petrosino, J. F., et al. (2020). SMRT Sequencing of Paramecium Bursaria Chlorella Virus-1 Reveals Diverse Methylation Stability in Adenines Targeted by Restriction Modification Systems. Frontiers in Microbiology 11. Available at: https://www.frontiersin.org/articles/10.3389/fmicb.2020.00887 [Accessed December 9, 2022].

De Vuyst, G., Aci, S., Genest, D., and Culard, F. (2005). Atypical Recognition of Particular DNA Sequences by the Archaeal Chromosomal MC1 Protein. Biochemistry 44, 10369–10377. doi: 10.1021/bi0474416.

Domingo, E., and Perales, C. (2019). Viral quasispecies. PLOS Genetics 15, e1008271. doi: 10.1371/journal.pgen.1008271.

Domingo, E., Sheldon, J., and Perales, C. (2012). Viral quasispecies evolution. Microbiol Mol Biol Rev 76, 159–216. doi: 10.1128/MMBR.05023-11.

Domingo, E., Soria, M. E., Gallego, I., de Ávila, A. I., García-Crespo, C., Martínez-González, B., et al. (2020). A new implication of quasispecies dynamics: Broad virus diversification in absence of external perturbations. Infection, Genetics and Evolution 82, 104278. doi: 10.1016/j.meegid.2020.104278.

Duffy, S. (2018). Why are RNA virus mutation rates so damn high? PLoS Biol 16, e3000003. doi: 10.1371/journal.pbio.3000003.

Etten, J., Dunigan, D., Nagasaki, K., Schroeder, D., Grimsley, N., Brussaard, C., et al. (2020). “Phycodnaviruses (Phycodnaviridae),” in Reference Module in Life Sciences doi: 10.1016/B978-0-12-809633-8.21291-0.

Filée, J. (2018). Giant viruses and their mobile genetic elements: the molecular symbiosis hypothesis. Current Opinion in Virology 33, 81–88. doi: 10.1016/j.coviro.2018.07.013.

Fischer, M. G. (2021). The Virophage Family Lavidaviridae. Current Issues in Molecular Biology 40, 1–24. doi: 10.21775/cimb.040.001.

Fischer, M. G., Kelly, I., Foster, L. J., and Suttle, C. A. (2014). The virion of Cafeteria roenbergensis virus (CroV) contains a complex suite of proteins for transcription and DNA repair. Virology 466–467, 82–94. doi: 10.1016/j.virol.2014.05.029.

Gallot-Lavallée, L., and Blanc, G. (2017). A Glimpse of Nucleo-Cytoplasmic Large DNA Virus Biodiversity through the Eukaryotic Genomics Window. Viruses 9, 17. doi: 10.3390/v9010017.

Gann, E. R., Truchon, A. R., Papoulis, S. E., Dyhrman, S. T., Gobler, C. J., and Wilhelm, S. W. (2022). Aureococcus anophagefferens (Pelagophyceae) genomes improve evaluation of nutrient acquisition strategies involved in brown tide dynamics. Journal of Phycology 58, 146–160. doi: 10.1111/jpy.13221.

Gastrich, M. D., Leigh-Bell, J. A., Gobler, C. J., Roger Anderson, O., Wilhelm, S. W., and Bryan, M. (2004). Viruses as potential regulators of regional brown tide blooms caused by the alga,Aureococcus anophagefferens. Estuaries 27, 112–119. doi: 10.1007/BF02803565.

Gobler, C., Anderson, O., Gastrich, M., and Wilhelm, S. (2007). Ecological aspects of viral infection and lysis in the harmful brown tide alga Aureococcus anophagefferens. Aquat. Microb. Ecol. 47, 25–36. doi: 10.3354/ame047025.

Gontero, B., and Maberly, S. C. (2012). An intrinsically disordered protein, CP12: jack of all trades and master of he Calvin cycle. Biochem Soc Trans 40, 995–999. doi: 10.1042/bst20120097.

Guillou, L., Bachar, D., Audic, S., Bass, D., Berney, C., Bittner, L., et al. (2013). The Protist Ribosomal Reference database (PR2): a catalog of unicellular eukaryote Small Sub-Unit rRNA sequences with curated taxonomy. Nucleic Acids Research 41, D597–D604. doi: 10.1093/nar/gks1160.

Hingamp, P., Grimsley, N., Acinas, S. G., Clerissi, C., Subirana, L., Poulain, J., et al. (2013). Exploring nucleo-cytoplasmic large DNA viruses in Tara Oceans microbial metagenomes. ISME J 7, 1678–1695. doi: 10.1038/ismej.2013.59.

Jeudy, S., Rigou, S., Alempic, J.-M., Claverie, J.-M., Abergel, C., and Legendre, M. (2020). The DNA methylation landscape of giant viruses. Nat Commun 11, 2657. doi: 10.1038/s41467-020-16414-2.

Jian, H., Yi, Y., Wang, J., Hao, Y., Zhang, M., Wang, S., et al. (2021). Diversity and distribution of viruses inhabiting the deepest ocean on Earth. ISME J 15, 3094–3110. doi: 10.1038/s41396-021-00994-y.

Jurasz, H., Pawłowski, T., and Perlejewski, K. (2021). Contamination Issue in Viral Metagenomics: Problems, Solutions, and Clinical Perspectives. Frontiers in Microbiology 12. Available at: https://www.frontiersin.org/articles/10.3389/fmicb.2021.745076 [Accessed December 9, 2022].

Kanehisa, M., Sato, Y., Kawashima, M., Furumichi, M., and Tanabe, M. (2016). KEGG as a reference resource for gene and protein annotation. Nucleic Acids Research 44, D457–D462. doi: 10.1093/nar/gkv1070.

Kapitonov, V. V., and Jurka, J. (2006). Self-synthesizing DNA transposons in eukaryotes. Proceedings of the National Academy of Sciences 103, 4540–4545. doi: 10.1073/pnas.0600833103.

Katoh, K., and Standley, D. M. (2013). MAFFT Multiple Sequence Alignment Software Version 7: Improvements in Performance and Usability. Mol Biol Evol 30, 772–780. doi: 10.1093/molbev/mst010.

Khalil, J. Y. B., Andreani, J., and La Scola, B. (2016). Updating strategies for isolating and discovering giant viruses. Current Opinion in Microbiology 31, 80–87. doi: 10.1016/j.mib.2016.03.004.

Koonin, E. V., and Krupovic, M. (2017). Polintons, virophages and transpovirons: a tangled web linking viruses, transposons and immunity. Current Opinion in Virology 25, 7–15. doi: 10.1016/j.coviro.2017.06.008.

Krasovec, M., Vancaester, E., Rombauts, S., Bucchini, F., Yau, S., Hemon, C., et al. (2018). Genome Analyses of the Microalga Picochlorum Provide Insights into the Evolution of Thermotolerance in the Green Lineage. Genome Biol Evol 10, 2347–2365. doi: 10.1093/gbe/evy167.

Krupovic, M., and Koonin, E. V. (2015). Polintons: a hotbed of eukaryotic virus, transposon and plasmid evolution. Nat Rev Microbiol 13, 105–115. doi: 10.1038/nrmicro3389.

La Scola, B., Desnues, C., Pagnier, I., Robert, C., Barrassi, L., Fournous, G., et al. (2008). The virophage as a unique parasite of the giant mimivirus. Nature 455, 100–104. doi: 10.1038/nature07218.

Lawrence, J., and Suttle, C. (2004). Effect of viral infection on sinking rates of Heterosigma akashiwo and its implications for bloom termination. Aquat. Microb. Ecol. 37, 1–7. doi: 10.3354/ame037001.

Luo, E., Eppley, J. M., Romano, A. E., Mende, D. R., and DeLong, E. F. (2020). Double-stranded DNA virioplankton dynamics and reproductive strategies in the oligotrophic open ocean water column. ISME J 14, 1304–1315. doi: 10.1038/s41396-020-0604-8.

Marshall, O. J. (2004). PerlPrimer: cross-platform, graphical primer design for standard, bisulphite and real-time PCR. Bioinformatics 20, 2471–2472. doi: 10.1093/bioinformatics/bth254.

Mikheenko, A., Saveliev, V., and Gurevich, A. (2016). MetaQUAST: evaluation of metagenome assemblies. Bioinformatics 32, 1088–1090. doi: 10.1093/bioinformatics/btv697.

Moniruzzaman, M., Weinheimer, A. R., Martinez-Gutierrez, C. A., and Aylward, F. O. (2020). Widespread endogenization of giant viruses shapes genomes of green algae. Nature 588, 141–145. doi: 10.1038/s41586-020-2924-2.

Nerlich, A., von Orlow, M., Rontein, D., Hanson, A. D., and Dörmann, P. (2007). Deficiency in phosphatidylserine decarboxylase activity in the psd1 psd2 psd3 triple mutant of Arabidopsis affects phosphatidylethanolamine accumulation in mitochondria. Plant Physiol 144, 904–914. doi: 10.1104/pp.107.095414.

Pagarete, A., Grébert, T., Stepanova, O., Sandaa, R.-A., and Bratbak, G. (2015). Tsv-N1: A Novel DNA Algal Virus that Infects Tetraselmis striata. Viruses 7, 3937–3953. doi: 10.3390/v7072806.

Palermo, C. N., Shea, D. W., and Short, S. M. (2021). Analysis of Different Size Fractions Provides a More Complete Perspective of Viral Diversity in a Freshwater Embayment. Applied and Environmental Microbiology 87, e00197–21. doi: 10.1128/AEM.00197-21.

Price, M. N., Dehal, P. S., and Arkin, A. P. (2009). FastTree: Computing Large Minimum Evolution Trees with Profiles instead of a Distance Matrix. Mol Biol Evol 26, 1641–1650. doi: 10.1093/molbev/msp077.

Redrejo-Rodríguez, M., and Salas, M. L. (2014). Repair of base damage and genome maintenance in the nucleo-cytoplasmic large DNA viruses. Virus Research 179, 12–25. doi: 10.1016/j.virusres.2013.10.017.

Roux, S., Chan, L.-K., Egan, R., Malmstrom, R. R., McMahon, K. D., and Sullivan, M. B. (2017). Ecogenomics of virophages and their giant virus hosts assessed through time series metagenomics. Nat Commun 8, 858. doi: 10.1038/s41467-017-01086-2.

Sheng, Y., Wu, Z., Xu, S., and Wang, Y. (2022). Isolation and Identification of a Large Green Alga Virus (Chlorella Virus XW01) of Mimiviridae and Its Virophage (Chlorella Virus Virophage SW01) by Using Unicellular Green Algal Cultures. J Virol 96, e0211421. doi: 10.1128/jvi.02114-21.

Short, S. M., Staniewski, M. A., Chaban, Y. V., Long, A. M., and Wang, D. (2020). Diversity of Viruses Infecting Eukaryotic Algae. Current Issues in Molecular Biology, 29–62. doi: 10.21775/cimb.039.029.

Smoot, M. E., Ono, K., Ruscheinski, J., Wang, P.-L., and Ideker, T. (2011). Cytoscape 2.8: new features for data integration and network visualization. Bioinformatics 27, 431–432. doi: 10.1093/bioinformatics/btq675.

Steinegger, M., and Söding, J. (2017). MMseqs2 enables sensitive protein sequence searching for the analysis of massive data sets. Nature Biotechnology 35, 1026–1028. doi: 10.1038/nbt.3988.

Suttle, C. A., and Wilhelm, S. W. (1999). Viruses and Nutrient Cycles in the Sea. 49, 8.

Tahlan, K., Park, H. U., Wong, A., Beatty, P. H., and Jensen, S. E. (2004). Two Sets of Paralogous Genes Encode the Enzymes Involved in the Early Stages of Clavulanic Acid and Clavam Metabolite Biosynthesis in Streptomyces clavuligerus. Antimicrob Agents Chemother 48, 930–939. doi: 10.1128/AAC.48.3.930-939.2004.

Tatusov, R. L., Galperin, M. Y., Natale, D. A., and Koonin, E. V. (2000). The COG database: a tool for genome-scale analysis of protein functions and evolution. Nucleic Acids Res 28, 33–36.

Untergasser, A., Cutcutache, I., Koressaar, T., Ye, J., Faircloth, B. C., Remm, M., et al. (2012). Primer3—new capabilities and interfaces. Nucleic Acids Res 40, e115–e115. doi: 10.1093/nar/gks596.

Van Etten, J. L., Burbank, D. E., Xia, Y., and Meints, R. H. (1983). Growth cycle of a virus, PBCV-1, that infects Chlorella-like algae. Virology 126, 117–125. doi: 10.1016/0042-6822(83)90466-X.

Wilson, W. H., Van Etten, J. L., and Allen, M. J. (2009). The Phycodnaviridae: The Story of How Tiny Giants Rule the World. Curr Top Microbiol Immunol 328, 1–42.

Yau, S., Caravello, G., Fonvieille, N., Desgranges, É., Moreau, H., and Grimsley, N. (2018). Rapidity of Genomic Adaptations to Prasinovirus Infection in a Marine Microalga. Viruses 10, 441. doi: 10.3390/v10080441.

Yau, S., Lauro, F. M., DeMaere, M. Z., Brown, M. V., Thomas, T., Raftery, M. J., et al. (2011). Virophage control of antarctic algal host-virus dynamics. Proceedings of the National Academy of Sciences 108, 6163–6168. doi: 10.1073/pnas.1018221108.

Yutin, N., Shevchenko, S., Kapitonov, V., Krupovic, M., and Koonin, E. V. (2015). A novel group of diverse Polinton-like viruses discovered by metagenome analysis. BMC Biology 13, 95. doi: 10.1186/s12915-015-0207-4.

Yutin, N., Wolf, Y. I., Raoult, D., and Koonin, E. V. (2009). Eukaryotic large nucleo-cytoplasmic DNA viruses: Clusters of orthologous genes and reconstruction of viral genome evolution. Virol J 6, 223. doi: 10.1186/1743-422X-6-223.

Zhang, W., Zhong, H., Lu, H., Zhang, Y., Deng, X., Huang, K., et al. (2018). Characterization of Ferredoxin-Dependent Biliverdin Reductase PCYA1 Reveals the Dual Function in Retrograde Bilin Biosynthesis and Interaction With Light-Dependent Protochlorophyllide Oxidoreductase LPOR in Chlamydomonas reinhardtii. Front Plant Sci 9, 676. doi: 10.3389/fpls.2018.00676.

